## Supplementary text and figures for "Complementary resource preferences spontaneously emerge in diauxic microbial communities"

This file contains:

Supplementary Figures, Fig. S1 to S6
Supplementary Text, sections A to I

#### Supplementary Figures

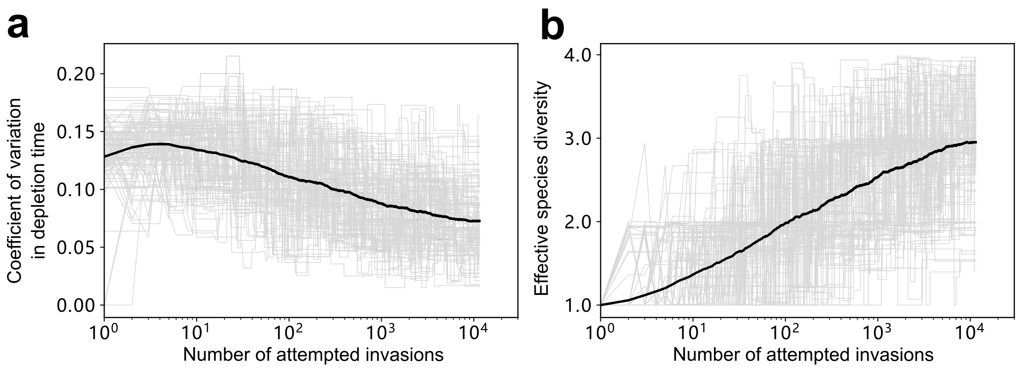

Figure S1: **Community-level properties during the assembly process.** Data from the same set of simulations as in Fig. 2. Solid lines in all subplots are averaged over all simulated communities at each invasion during the assembly process. Grey lines correspond to 100 random individual community assembly simulations. **(a)** Coefficient of variation in $T_i$'s slowly decreases during the assembly process. **(b)** Effective diversity, here defined by inverse Simpson's index, increases during the assembly.

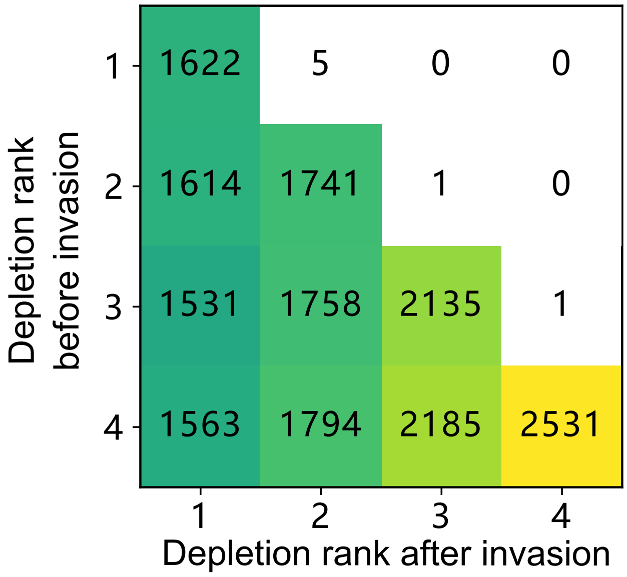

Figure S2: **Depletion order of the invader's top choice before and after the invasion.** Data from the same set of simulations as in Fig. 2. To eliminate the unnecessary randomness in the early stage of assembly, successful invasions before the 100th attempt were not counted. In each grid there displays the count of the corresponding event.

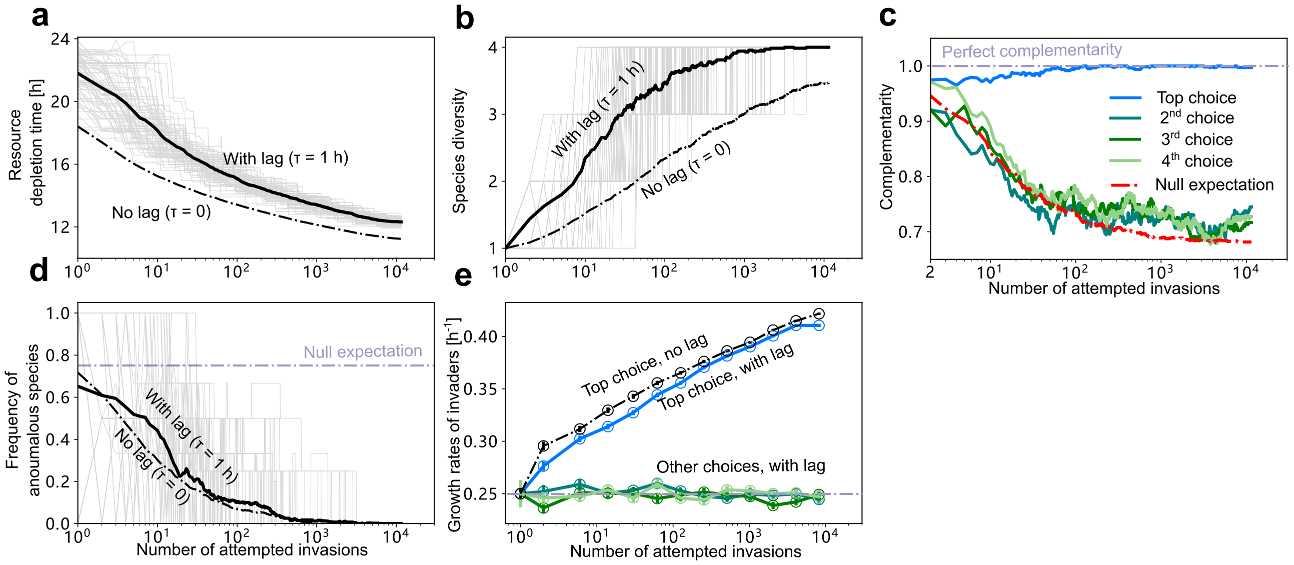

Figure S3: **Effects of a finite lag on community assembly** Both "with lag" and"no lag" data come from the same set of simulations as in Fig. 5. In (a), (b) and (d) the grey lines display the data from the 100 simulations where the lag time is 1 hr.

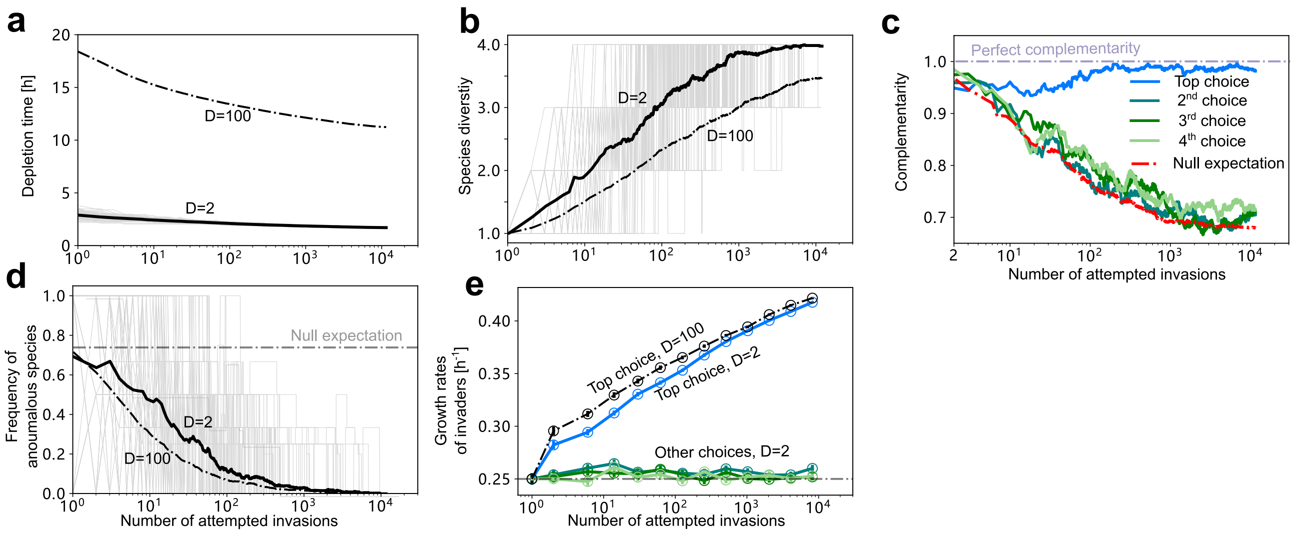

Figure S4: **Effects of dilution factor being small on community assembly** Data of $D = 100$ comes from the same set of simulations as in Fig. 2. The dilution factor $D$ is the only modified parameter when simulating the $D = 2$ data. In (a), (b) and (d) the grey lines display the $D = 2$ data, which contains 100 simulations of independent community assembly.

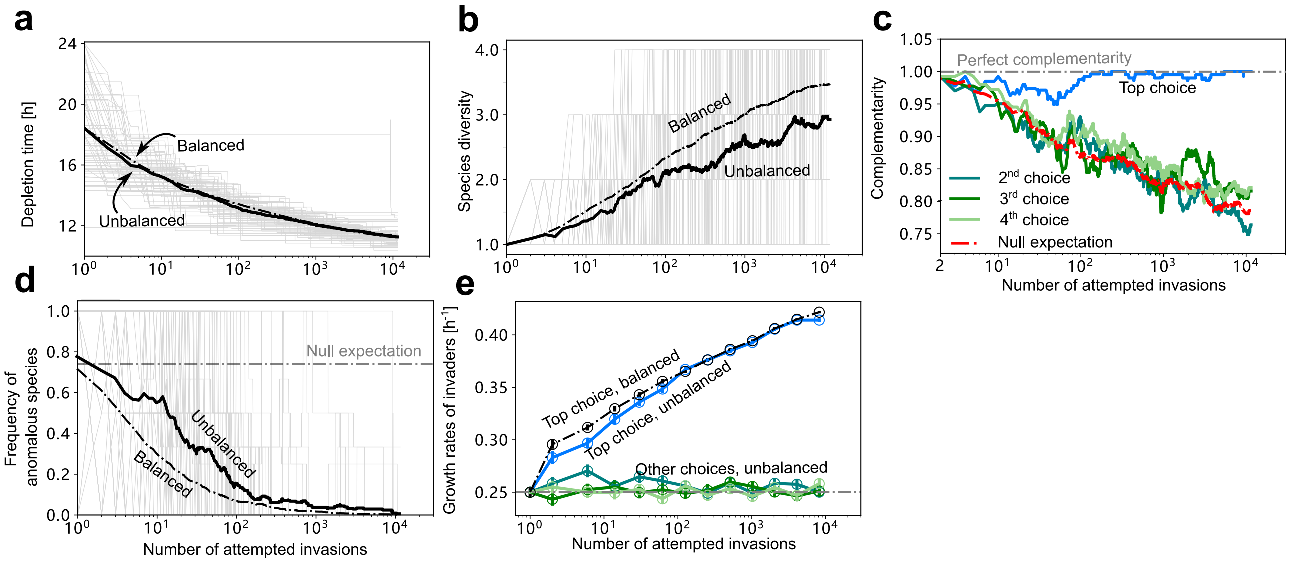

Figure S5: **Effects of extremely unbalanced nutrient supply on community assembly** Data of balanced resources comes from the same set of simulations as in Fig. 2. As for the unbalanced resources, the 4 resources were set to a ratio of 100:1:1:1 at the beginning of each dilution cycle, with the sum of all resource concentrations kept at the same level as in the balanced case. In (a), (b), and (d) the grey lines display the data of unbalanced resources, which contains 100 simulations of independent community assembly.

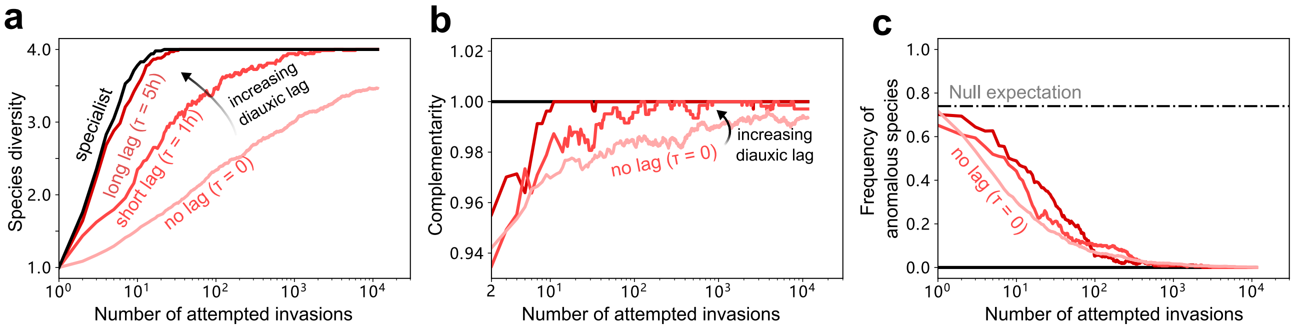

Figure S6: **Compare diauxic shifters with lags to specialists** All diauxic shifters' data come from the same set of simulations as in Fig. 5. The data of the specialists were simulated in the following manner. We generated the top choice growth rates of those specialists from the same distribution ($N(0.5, 0.1^2)$ rectified at 0.1). We then introduced these species one by one by the following rule: after each invasion, on each resource, the specialists with the highest growth rates win and stay. This includes 100 independent community assembly processes. In all three panels, the black lines correspond to the specialists, the red lines display the data from lag time 5 hr, the orange lines show the case of lag time 1 hr, and the brown lines correspond to simulations without lag.

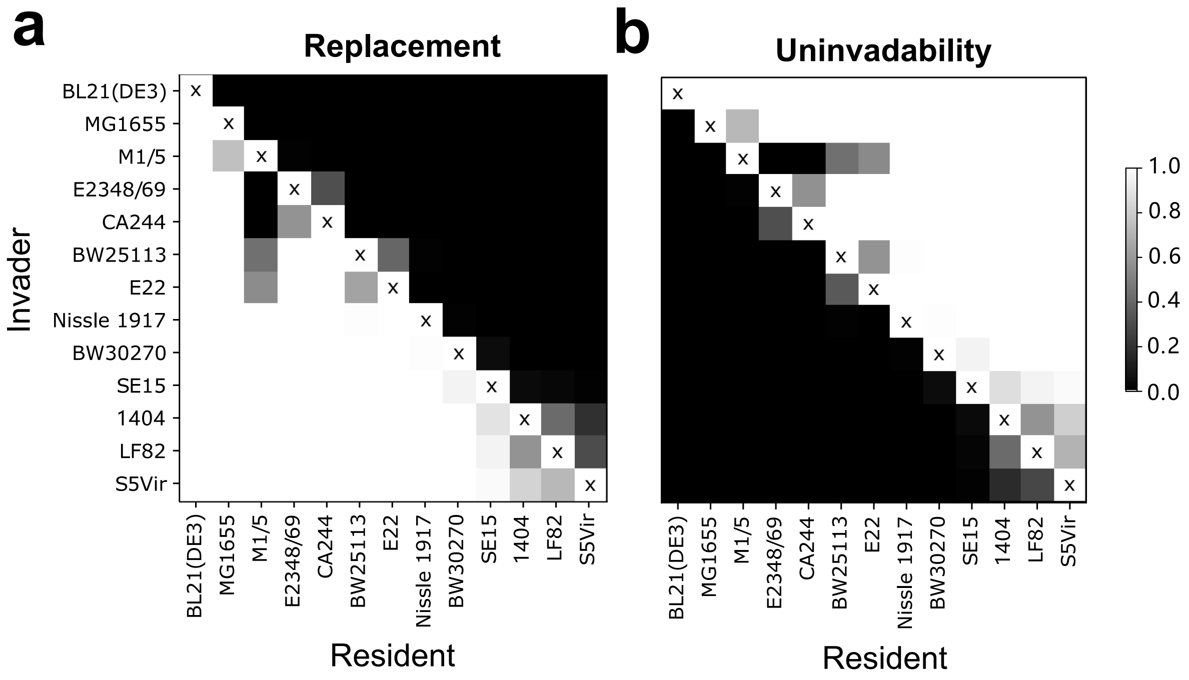

Figure S7: **Numerical invasion experiments of *E. coli* strains studied in** [1]**.** A strain on the $y$-axis invades the monoculture of a strain on the $x$-axis, and the greyscale represents the frequency of both strains survive in the steady state, counted from 100 runs where lag time values were individually sampled. In **(a)** shows the frequency where the invader replaces the resident, and in **(b)** shows the frequency where the invader does not survive in the end.

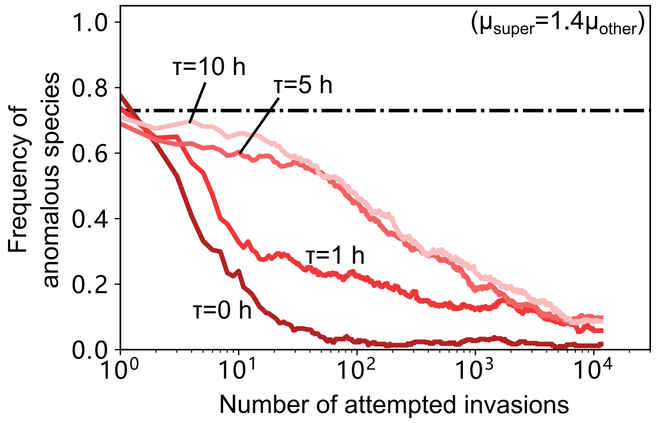

Figure 8: **Effects of having unbalanced growth rate distributions on different resources** Growth rate distribution on a "super" resource $R_{super}$ is set at $N(\mu_{super} = 0.35, 0.05^2)$, while such distribution on the rest 3 resources remain at $N(\mu_{other} = 0.25, 0.05^2)$. All other parameters are the same as the simulations regarding Fig. 2. Lags promote the survival of anomalous species when a "super" resource is present. The black dash-dotted line shows the proportion of anomalous species in the species pool. Lines of shades of red display frequency of anomalous species during the community assembly, averaged over 100 community assembly simulations.

**Supplementary Text**
**Complementary resource preferences spontaneously emerge in diauxic microbial communities**

**I Representing the co-utilizing microbes in the time plane 35**

**J Coexistence of the diauxic species having the same preferential orders 37**

**K What happens to complementarity and diversity when the growth rate distributions are not similar for different sources 40**

### A Depletion order and times in the light of our Tilman-inspired approach

In a standard serial dilution experiment, resources get resupplied in the same fixed amounts at the beginning of each dilution cycle. The microbial community populating the environment comes to each new step diluted $D-$fold, compared to the values of the species concentrations acquired at the end of the previous dilution cycle. $D$ here is the dilution coefficient.

Depletion order is the order in which resources disappear from the environment one by one during one given dilution cycle, getting consumed by the microbes.

Consider first a single microbe invading an abiotic medium containing two resources in the concentrations $R_1$ and $R_2$ (by $R$ capital we designate both resources and their concentration values, depending on the context). Suppose $R_1$ precedes $R_2$ in the microbe's personal preferential list. If the bacterium invades in a seed amount, it may not be able to consume even $R_1$ during the first dilution cycle. If that is the case, during the first cycle it does not even touch $R_2$. However, if the cycle lasts long enough, the microbe grows more than $D-$fold during the step, so that it comes to the next dilution step in greater concentration. Eventually it gets fast enough to consume both $R_1$ and $R_2$, one by one in a diauxie manner, in the order determined by its own list of preferences. When it reaches the point of growing precisely $D-$fold during a dilution step, it comes to every next cycle with the same initial concentration as to the previous one, and thus, in a sense, a steady state arises. Let $T_1$ be the total lifetime of the resource $R_1$ and $T_2$ that of $R_2$ during a dilution cycle. Obviously, as $R_1$ gets consumed first, $T_2 > T_1$, and $T_2$ ends up to be the effective time of the microbe growth during the dilution cycle.

For the steady state, we write:

$$N_\alpha \exp\Big(g_{\alpha 1}T_1 + g_{\alpha 2}\left(T_2 - T_1\right)\Big) = DN_\alpha, \tag{S.1}$$

so that

$$g_{\alpha 1}T_1 + g_{\alpha 2}\left(T_2 - T_1\right) = \log\, D, \tag{S.2}$$

where $N_\alpha$ is the initial steady state concentration of the bacterial species $B_\alpha$, $g_{\alpha i}$ is its growth rate on the resource number $i$, time $T_1$ the bacterium spends growing exponentially on the resource $R_1$ and for the rest $T_2 - T_1$ it grows on the resource $R_2$.

The steady state concentration $N_\alpha$ and the times $T_1$, $T_2$ are determined by the initial (for each cycle) resource concentrations $R_1$ and $R_2$ in the following way. First, starting a new dilution cycle with the initial steady state concentration $N_\alpha$, the microbe ends it having consumed both $R_1$ and $R_2$, so that

$$DN_\alpha = N_\alpha + R_1 + R_2, \tag{S.3}$$

or else

$$N_\alpha = \frac{R_1 + R_2}{D - 1}, \tag{S.4}$$

note that in the above we do not take yields into account.

As the resource $R_1$ gets consumed in the time period $T_1$ (the depletion time of $R_1$)

$$N_\alpha\, \exp\left(g_{\alpha 1}T_1\right) = N_\alpha + R_1, \tag{S.5}$$

which, combined with $(S.4)$, gives

$$T_1 = \frac{1}{g_{\alpha 1}}\, \log\, \Big(\frac{DR_1 + R_2}{R_1 + R_2}\Big). \tag{S.6}$$

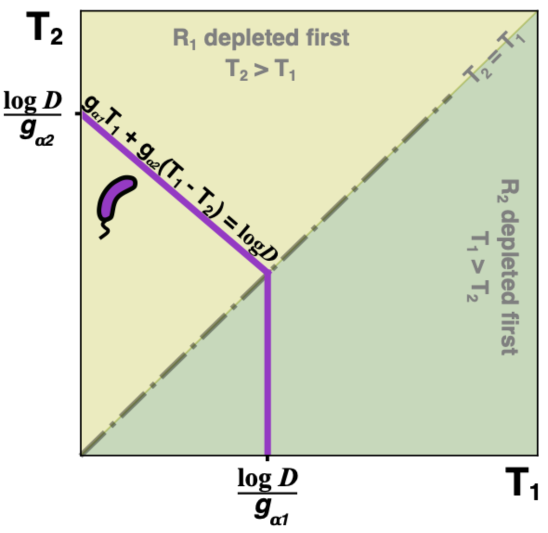

Figure S9: **Tilman-inspired diagram for one microbe.** Zero Net Growth Isocline (ZNGI) of a single microbe $B_\alpha$ growing in an environment that contains two sources $R_1$ and $R_2$ with the growth rates $g_{\alpha 1}$ and $g_{\alpha 2}$ correspondingly. The environment is subject to serial dilution with the dilution coefficient $D$. The axes $T_1$, $T_2$ represent the corresponding depletion times of $R_1$, $R_2$. In the given environment, the microbe's top choice is $R_1$. If the latter is depleted last (the lower triangle of the diagram, $T_1 > T_2$), the microbe grows on it with the growth rate $g_{\alpha 1}$, never exhibiting a diauxic shift.

In the similar way, for the rest of the effective growth time $T_2 - T_1$, $R_2$ is consumed, and we get:

$$T_2 - T_1 = \frac{1}{g_{\alpha 2}} \log \left( \frac{D(R_1 + R_2)}{DR_1 + R_2} \right). \tag{S.7}$$

In the case when a single microbe resides in the environment containing two resources, the depletion order is defined by the microbe's preferences, and the steady state concentration of the microbe as well as the depletion times depend on the initial resource concentrations of each dilution cycle.

When a single microbe invades a multi-nutrient environment, the resources concentration values being $R_1, R_2, ..., R_k, ..., R_n$, the sequence in the above reflecting the preferential order of that particular microbe $B_\alpha$, the depletion times are obtained in the way similar to the above, so that

$$\Delta T_k = T_k - T_{k-1} = \frac{1}{g_{\alpha k}} \log \left( \frac{D(R_1 + R_2 + ... + R_{k-1} + R_k) + R_{k+1} + ... + R_n}{D(R_1 + R_2 + ... + R_{k-1}) + R_k + R_{k+1} + ... + R_n} \right) \tag{S.8}$$

In the case of a balanced nutrient supply $R_1 = R_2 = ... = R_n$ and $D \gg n$ one can write

$$\Delta T_1 = T_1 \simeq \frac{1}{g_{\alpha 1}} \log \left( \frac{D}{n} \right), \; \Delta T_2 = T_2 - T_1 \simeq \frac{1}{g_{\alpha 2}} \log \frac{2}{1}, \; \Delta T_3 = T_3 - T_2 \simeq \frac{1}{g_{\alpha 3}} \log \frac{3}{2} ... \tag{S.9}$$

so that $\Delta T_k = (1/g_{\alpha k}) \log(n/(n-1))$. So, at large values of the dilution coefficient $D$ the microbe spends a huge share of the growing time consuming its top-choice resource. Its concentration gets multiplied by approximately $D/n$ upon using up $R_1$, while consummation of $R_2$ just doubles it, and so on.

For a medium containing two resources $R_1$ and $R_2$, invaded by a single microbe $B_\alpha$, we could construct a Tilman-like diagram (see Fig. S9). We have found that, for the serial dilution setting, the state-defining role of the resource concentrations assigned to the axes on the original Tilman's plane[2] should be played instead by the depletion times $T_1$ and $T_2$. Any point $(T_1, T_2)$ of the line described by the equation (S.2) represents the pair of depletion times in a steady state, determined by a particular set of the initial resource concentrations. The line is only valid when $T_2 > T_1$ — that would be the depletion order determined by the nutritional preferences of the microbe $B_\alpha$.

When yet another microbe $B_\beta$ tries to invade in a seed amount, the challenge is to survive and grow in the medium shaped for it by the microbe $B_\alpha$. During the first dilution cycle at which $B_\beta$ is present, as it has been pointed out in the main text, its influence on the resources lifetimes is negligible. Thus

$$g_{\beta 1}\, T_1 + g_{\beta 2}\, (T_2 - T_1) > \log D \tag{S.10}$$

$(g_{\beta 1}, g_{\beta 2}$ being the new invader's growth rates on the resources $R_1$ and $R_2)$ is the criterion of whether $B_\beta$ is able to grow in the invaded medium **if the resources' ranks in its preferential list correspond to the depletion order of the resources in the invaded environment.**

For a microbe $B_\gamma$ that prefers $R_2$ to $R_1$, the criterion of successful invasion of the environment shaped by $B_\alpha$ would be

$$g_{\gamma 2}\, T_2 > \log D \tag{S.11}$$

meaning that what matters is whether it can grow on its top-choice resource (of those present in the environment) fast enough.

The Tilman-inspired diagrams with the sole-microbe steady state ZNGI lines of $B_\beta$ and $B_\gamma$, together with that of $B_\alpha$, are featured in Fig.S10. Each of them corresponds to the set of the steady states realizable under different nutrition conditions (i. e., the ratio of the concentrations $R_1/R_2$) when one microbe ($B_\alpha$, $B_\beta$ or $B_\gamma$) invades the abiotic environment.

In the usual way, the approach invented by Tilman enables one to determine whether a new-coming invader will survive under given circumstances. If the given steady state depletion times $T_1$ and $T_2$ define a point situated under the steady state lifeline (the ZNGI, to use Tilman's original terminology) of the invader, then the latter would not be able to grow. If, on the other hand, the depletion times characterizing the steady state map themselves to the point above the ZNGI of the invader, the latter will start growing and reshaping the environment until the depletion times set somewhere on its own ZNGI.

The numbers $I$, $II$, $III$ in Fig.S10 mark points in the Time-plane at which the coexistence of two microbes is possible. In absence of strong trade-off conditions there are, in general, no points at which three or more lines intersect, so that the number of survivors in a steady state can never exceed the number of nutrition sources. In presence of all the three contestants, the only possible coexistence point is $II$, at which the microbe $B_\alpha$ preferring $R_1$ to $R_2$ coexists with $B_\gamma$ with the reversed preference order: the top choice $R_2$, the second choice $R_1$. In presence of the species $B_\gamma$ the species $B_\alpha$ and $B_\beta$ cannot coexist (at point $I$) as the diagram shows that, at the point $I$, $B_\gamma$ can grow. The coexistence point $III$ of the pair $B_\alpha$, $B_\gamma$, on the other hand, is disabled by the presence of the species $B_\beta$ whose ZNGI goes lower.

For the two species $B_\alpha$ and $B_\beta$ to be able to coexist, we must have

$$\begin{cases} g_{\alpha 1}\, T_1 + g_{\alpha 2}\, (T_2 - T_1) = \log D \\ g_{\beta 1}\, T_1 + g_{\beta 2}\, (T_2 - T_1) = \log D \end{cases} \tag{S.12}$$

From that we get the depletion times that permit coexistence of $B_\alpha$ and $B_\beta$ in absence of $B_\gamma$ (point $I$ in Fig.S10)

$$\begin{cases} T_1 = \dfrac{g_{\beta 2} - g_{\alpha 2}}{g_{\alpha 1}\, g_{\beta 2} - g_{\beta 1}\, g_{\alpha 2}} \log D \\ \\ T_2 = \dfrac{g_{\alpha 1} - g_{\alpha 2} - g_{\beta 1} + g_{\beta 2}}{g_{\alpha 1}\, g_{\beta 2} - g_{\alpha 2}\, g_{\beta 1}} \log D. \end{cases} \tag{S.13}$$

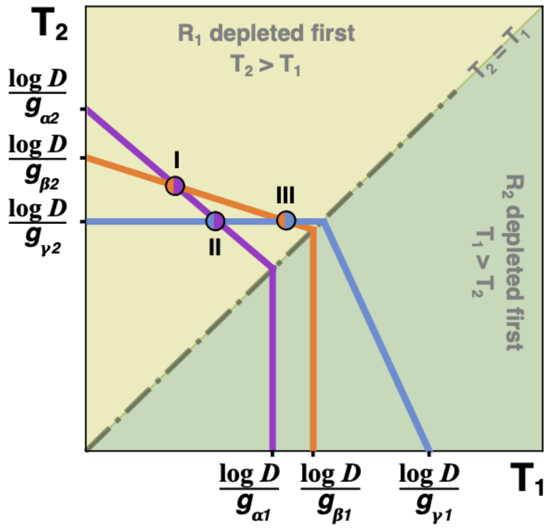

Figure S10: **Tilman-inspired diagrams featuring possible coexistence of microbes with similar and different nutrition preferences.** The ZNGI lines of the microbes $B_\alpha$ (lilac) and $B_\beta$ (orange) with the same top choice $R_1$ intersect at the point **I**, at which both can coexist within a certain range of the $R_1/R_2$ ratio. At that point however $B_\gamma$ (light blue) with the top choice $R_2$ can invade, supplanting $B_\beta$ and bringing the system to the coexisting point **II**. In absence of $B_\alpha$ the microbes $B_\beta$ and $B_\gamma$ are able to coexist at the point **III**.

To get the coexistence conditions for $B_\alpha$ and $B_\gamma$ sharing the same environment (point II), as the coexistence happens at $T_2 > T_1$ and thus $B_\gamma$ uses only its top choice $R_2$ through the whole growing period, we write

$$\begin{cases} g_{\alpha 1}\, T_1 + g_{\alpha 2}\,(T_2 - T_1) = \log D \\ g_{\gamma 2}\, T_2 = \log D \end{cases} \tag{S.14}$$

That leads to

$$\begin{cases} T_1 = \dfrac{g_{\alpha 2} - g_{\gamma 2}}{g_{\gamma 2}\,(g_{\alpha 1} - g_{\alpha 2})} \log D \\ \\ T_2 = \dfrac{\log D}{g_{\gamma 2}} \end{cases} \tag{S.15}$$

The coexistence point $III$ at which $B_\beta$ and $B_\gamma$ can share the same environment in absence of the microbe $B_\alpha$ is defined by the formulae similar to (S.15), when obvious substitutions are made.

The conditions (S.13) and (S.15) show that the depletion times enabling coexistence (in the case of maximum diversity, when the number of microbes equals that of the sources of nutrition) are fully determined by the relevant growth rates. That kind of coexistence required the ratio of the resource concentrations $R_1/R_2$ to fall within a certain range determined by the growth rates and the dilution coefficient.

The steady state condition imposed on the initial (at the beginning of each dilution step) microbe concentrations in the case of $B_\alpha$ and $B_\beta$ peacefully coexisting:

$$DN_\alpha + DN_\beta = R_1 + R_2 + N_\alpha + N_\beta \tag{S.16}$$

It $T_1$ is the depletion time of $R_1$ from (S.13), in the steady state in which $B_\alpha$ coexists with $B_\beta$ one must have

$$N_\alpha \exp\left(g_{\alpha 1}\, T_1\right) + N_\beta \exp\left(g_{\beta 1}\, T_1\right) = R_1 + N_\alpha + N_\beta \tag{S.17}$$

Combining (S.16) and (S.13) to find steady state concentration values of $B_\alpha$ and $B_\beta$ and requiring those to be positive, we get the condition

$$\frac{\exp\left(g_{\beta 1}\, T_1\right) - 1}{D - \exp\left(g_{\beta 1}\, T_1\right)} < \frac{R_1}{R_2} < \frac{\exp\left(g_{\alpha 1}\, T_1\right) - 1}{D - \exp\left(g_{\alpha 1}\, T_1\right)} \tag{S.18}$$

In Eq. (S.13) and (S.18) we implied that $g_{\alpha 1} > g_{\beta 1}$ and $g_{\alpha 2} < g_{\beta 2}$, as the diagram in the Fig.S10 would have it. This is the usual counter-ordered arrangement for the coexistence of the species with similar preferences.

In the case of $B_\alpha$ and $B_\gamma$ coexisting, their top choices being different, Eq. (S.17) relating the microbe and resource concentration values at the moment when $R_1$ (which is the first to disappear) is consumed, takes the form

$$N_\alpha \exp\left(g_{\alpha 1} T_1\right) = R_1 + N_\alpha \tag{S.19}$$

as the microbe $B_\beta$ never tastes $R_1$. Obtaining the condition for $B_\beta$ to be present in the steady state and thus have positive concentration value, we get simply:

$$\frac{R_1}{R_2} < \frac{\exp\left(g_{\alpha 1} T_1\right) - 1}{D - \exp\left(g_{\alpha 1} T_1\right)} \tag{S.20}$$

Comparing Eq. (S.18) with Eq. (S.20) we conclude that, generally speaking, the range of the ratio $R_1/R_2$ values favoring coexistence is larger when the microbes in question have complementary preferences.

Note that in the above case **the concentrations** $(R_1, R_2)$ **do not map point to point onto the Time-plane**. In the case of the coexistence of two microbes, be it the species with the similar or different nutrition preferences, a single point $(T_1, T_2)$ corresponds to a continuous interval of the ratio $R_1/R_2$, or, in the other words, to a band with non-parallel straight boundaries in the original Tilman's resource plane.

**The depletion order** in the case of two resources present in the environment, barring special cases, is defined by the absolute fastest consumer of its top-choice resource. Indeed, if the top-choice of such a fastest-growing microbe were the last to disappear, the champion microbe would grow on it through the whole dilution cycle and thus its concentration would acquire greater multiplication coefficient than that of any other competing microbe. So it will be the absolute winner — yet, in this case, it will shape the environment in such a way that its top-choice resource is the first to disappear. For the same reason, when the resources present in the environment are multiple, the absolute fastest grower will not have its top-choice resource to be the last to disappear, again barring certain special circumstances. In the multi-resource environment, however, the champion species cannot be expected to establish the depletion order all on its own.

Now to the special circumstances from the previous passage. Those have to do with the special species, in the main text referred to as **anomalous** (as opposed to **normal**) microbes. In the case of an environment containing just two resources, the anomalous microbes are those that grow faster on their second choice resource, and not on their top choice one. Such microbes can display a bizarre nutrition behaviour, as can be seen in the Fig. S11.

An anomalous microbe is easily distinguished on a Tilman-inspired diagram by the different slope, rising up from the relevant time-axis instead of falling to it, as is the case with the normal microbes' ZNGI. It turns out that even being the absolute fastest grower (but on its second choice!), such a species may not be able to invade an environment if its top choice (on which it grows slower) is the last to disappear. In presence of an anomalous species, bistability may arise. In the case shown in Fig. S11, the resulting steady state will depend on which species, the anomalous one or its normal counterpart, is the first to invade. The anomalous species can make life in a biocontainer less dull by setting off the predetermination; more of that in the section dedicated specially to the anomalous microbes and the reason of them being rare.

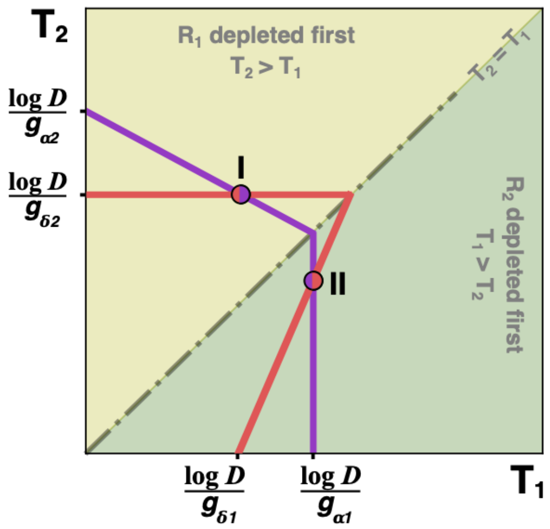

Figure S11: **An anomalous microbe competing with a normal one, represented by a Tilman-inspired diagram.** The anomalous microbe $B_\delta$ with the top choice $R_2$ (red ZNGI, with the positive slope at $T_1 > T_2$ displaying the abnormality) can coexist with the normal microbe $B_\alpha$ (lilac ZNGI, top choice $R_1$) at the points **I** and **II**. The coexistence at **I** is stable, while the coexistence at the point **II** is unstable. If prepared, the latter decays either down to the state at which $B_\delta$ (or $B_\alpha$) is the single resident, or up to the point of the stable coexistence **I**, depending on the value of the ratio $R_1/R_2$. Within a certain region of the values of $R_1/R_2$ bistability can occur. If $B_\alpha$ is the first to invade, it may either withstand the invasion by $B_\delta$ or let it share and coexist, depending on the value of $R_1/R_2$. If $B_\delta$ is the first to invade, it may shape the environment so that $B_\alpha$ cannon invade in its turn. Thus, depending on the order of invasion, the bistability between two different final steady states may be possible. Further investigation shows that, depending on the dilution coefficient and the growth rate parameters, the bistability may occur either between two monoculture states or between two states one of which is that of monoculture and the other is that of coexistence.

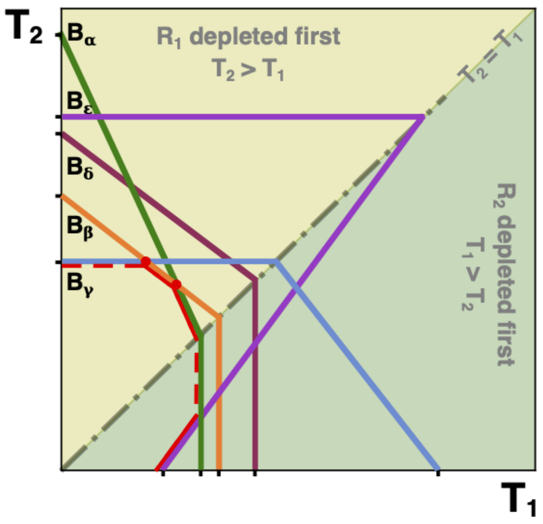

Figure S12: **The Pareto-optimal boundary.** Of all the competing microbes, each one except $B_\delta$ (auburn ZNGI, top choice $R_1$) participates in the forming of the Pareto-optimal boundary (marked in red). The parts of the POB marked by the dashed red line do not contain any points that correspond to the realizable steady states of the system. They are useful however to determine whether, for instance, $B_\gamma$ (light blue ZNGI, top choice $R_2$) can invade an environment shaped by, say, $B_\beta$ (orange ZNGI, top choice $R_1$) at a given steady state. When it can, the coexistence with $B_\beta$ will occur. As the point of coexistence belongs to the Pareto-optimal boundary, such a state will be uninvadable. At the balanced nutrient supply, the steady state of the system will most likely correspond to a point on the POB not far from the diagonal $T_1 = T_2$.

### B Pareto-optimal boundary and the one-microbe line

#### B.1 Relation to Tilman's graphical method

The original Tilman's diagrams that have resource concentrations for the axes enable one to predict which steady states can be uninvadable by visual means[3, 4]. For that, two things must appear on the diagram: first, the full set of the life vs death boundaries (ZNGI) for each species, to determine which species can grow when the others would die out; second, the so-called one-microbe lines. Each of the latter is the dynamic trajectory of a single-species state, traced from a particular abiotic point to that same species' zero net growth rate isoclines; at the intersection, a steady state arises. Usually the ZNGI lines are fully determined by the properties of the species they correspond to (e. g. growth rates, yields, maintenance costs etc.) and the dilution coefficient when relevant, while the one-microbe lines depend on the properties of a particular species as much as on the external conditions.

Our Tilman-inspired diagrams are a no less powerful instrument in this sense, to be handled with care, as, in general, they do not map to a resource plane (or space) point by point. In the serial dilution case, the ZNGI lines get scaled by the dilution coefficient $D$; otherwise, they are completely determined by the growth rates of the corresponding species.

#### B.2 The Pareto-optimal boundary

When there are many competitors in the environment, some of the species never get to survive, and some others can win the game only at exotic external conditions (e. g. large imbalance in the regular nutrient supply). The question of which of the species can boast a range of external conditions favorable to their survival at the expense of the other competitors is related to the notion of *Pareto-optimal boundary.*

A ZNGI line for a microbe $B_\alpha$ that prefers the resource $R_1$ to $R_2$ has the equation

$$g_{\alpha 1} T_1 + g_{\alpha 2}(T_2 - T_1) = \log\, D, \tag{S.21}$$

in the upper segment of the positive quarter of the Time plane $T_1, T_2$: here $T_2 > T_1$ so that $B_\alpha$ starts with $R_1$, finishes it till the end of the time period $T_1$ and then, for the time period of the length $T_2 - T_1$ grows on the resource $R_2$. Whenever it grows more than $D-$fold through a dilution cycle (i. e. the time point of the environment is positioned above that boundary), it survives; if however the times of the environment correspond to a point below the line (S.21), the species $B_\alpha$ is doomed to get scarce and vanish from the serially diluted environment. In the lower segment of the positive quarter of the Time plane we have $T_1 > T_2$, so that through the whole effective growth time of the dilution cycle $R_1$ is present and $B_\alpha$ does not have to switch eventually to $R_2$. The segment of the life vs death boundary for $B_\alpha$ belonging to this area has the equation

$$g_{\alpha 1} T_1 = \log\, D. \tag{S.22}$$

The latter segment does not account for any single-microbe steady state point due to the fact that, when single, $B_\alpha$ will eventually shape the environment in such a way that $R_1$ is depleted first. Still, it is useful to judge whether $B_\alpha$ can grow when it invades the environment in which $R_1$ is the last to be depleted, and whether it can possibly coexist with another microbe in such an environment.

In Fig. S12 five microbes $B_\alpha$, $B_\beta$, $B_\gamma$, $B_\delta$, $B_\epsilon$ compete for the resources $R_1$, $R_2$. The shapes of their ZNGI give one a hint that $B_\alpha$, $B_\beta$, $B_\delta$ prefer $R_1$ to $R_2$ while $B_\gamma$ and $B_\epsilon$ choose otherwise. It is also clear that $B_\epsilon$ is anomalous, meaning that when given a choice, it starts consuming $R_2$ rather than $R_1$ despite the fact that it grows faster on $R_1$. (It is useful to remember that the ZNGI lines hit the axes $T_1$ and $T_2$ at the points $(\log D)/g_1$, $(\log D)/g_2$ respectfully, so that the closer the point is to origin, the larger the corresponding growth rate.) **The lower envelope** of the zero net growth isoclines is the line indicating the winners of the game (either single species or coexisting pairs) under various external conditions. We refer to it as the **Pareto-optimal boundary**. In Fig. S12 it is drawn in red. The dashed segments of the Pareto-optimal boundary trace the depletion times that effectively cannot be realized in a steady state; any steady state which is uninvadable can be represented by a point on the solid red line.

##### B.3 One-microbe line

Now to the notion of the one-species or the one-microbe line. While in the models considered by Tilman[3, 4] the resources are being continuously depleted, and for that reason the trajectories in the resource plane are continuous, with the depletion times in the serial dilution experiment that cannot be the case. If, say, $B_\alpha$ is introduced to an abiotic environment of a serial dilution experiment with the dilution coefficient $D$ and initial nutrient abundances $R_1$, $R_2$ in a seed amount, it may not be able to consume the available amount of $R_1$ through the first dilution cycle, or even a couple of those. While this is the case, the depletion times remain $T_1 = T_2 = T$, where $T$ is the fixed durance of the dilution cycle, and the "one-microbe line" is reduced to a single point. When $B_\alpha$ is finally able to consume $R_1$, it may or may not manage to deplete $R_2$ during the cycle; if the latter is the case, $T_1$ diminishes, while $T_2$ remains equal to the durance of the cycle, so the point representing the state of the system in terms of the depletion times travels left along the line $T_2 = T$. When $B_\alpha$, finally, succeeds at depleting both $R_1$ and $R_2$ in one dilution cycle, we would have, for some dilution cycle number $k$:

$$B_\alpha^{(k)}\, \exp\left(g_{\alpha 1} T_1^{(k)}\right) = R_1 + B_\alpha^{(k)} \tag{S.23}$$

and

$$B_\alpha^{(k)}\, \exp\left(g_{\alpha 1} T_1^{(k)} + g_{\alpha 2}\left(T_2^{(k)} - T_1^{(k)}\right)\right) = R_1 + R_2 + B_\alpha^{(k)}, \tag{S.24}$$

where the upper index $(k)$ means that value relevant for the $k-$th dilution cycle.

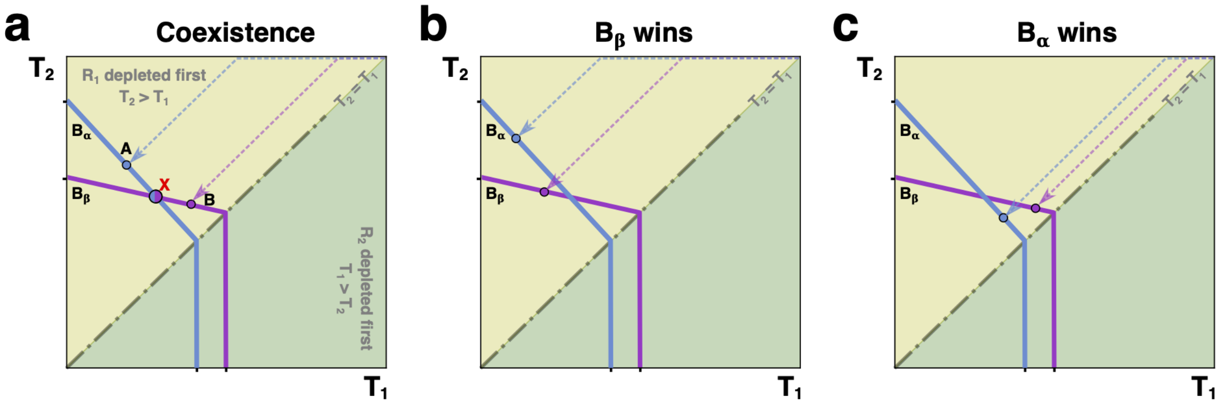

Figure S13: **The one-microbe lines indicating the winner(s).** **(a)** The (light blue, dotted) one-microbe line of the microbe $B_\alpha$ lands on the (light blue, solid) ZNGI of that same microbe at the point **A**. At this point, the species $B_\beta$ can invade. The (lilac, dotted) one-microbe line of the bacterium $B_\beta$ lands on the (lilac, solid) ZNGI of that species at the point **B**. At this point, the competing microbe $B_\alpha$ can invade. As a result of both invasions, the species $B_\alpha$ and $B_\beta$ will coexist at the point **X**. **(b)** The (light blue, dotted) one-microbe line of $B_\alpha$ lands on the (light blue, solid) ZNGI of that same species at the point where $B_\beta$ can invade. The (lilac, dotted) one-microbe line of $B_\beta$ lands on the species' ZNGI (lilac, solid) at the point where $B_\alpha$ cannot invade. Thus $B_\beta$ wins the competition. **(c)** The (light blue, dotted) one-microbe line of $B_\alpha$ lands on the species' ZNGI (light blue, solid) at the point where $B_\beta$ cannot invade. The (lilac, dotted) one-microbe line of $B_\beta$ lands on the (lilac, solid) ZNGI of that same species at the point where $B_\alpha$ can successfully invade. In this case, $B_\alpha$ is the sole winner.

Excluding $B_\alpha^{(k)}$ from both (S.23) and (S.24), we get

$$T_2^{(k)} = T_1^{(k)} + \frac{1}{g_{\alpha 2}} \log \left( \frac{R_2}{R_1} \left(1 - \exp\left(-g_{\alpha 1} T_1^{(k)}\right)\right) + 1 \right) \tag{S.25}$$

At $g_{\alpha 1} T_1^{(k)} \gg 1$ which is certainly true for reasonably balanced nutrient supply and $D \gg 1$, through the whole dynamics from that point on we would get $T_2 \sim T_1 + \text{const}$, so that the one-microbe line would go almost parallel to the diagonal $T_1 = T_2$. It is not continuous, so in reality the one-microbe line of $B_\alpha$ is a set of points belonging to the curve from (S.25). At the steady state $g_{\alpha 1} T_1 + g_{\alpha 2}(T_2 - T_1) = \log D$, so that using (S.25) we get for the steady state depletion times:

$$T_1 = \frac{1}{g_{\alpha 1}} \log \left( \frac{D R_1 + R_2}{R_1 + R_2} \right), \; T_2 = T_1 + \frac{1}{g_{\alpha 2}} \log \left( \frac{D\,(R_1 + R_2)}{D R_1 + R_2} \right), \tag{S.26}$$

same as we obtained in the previous sections.

In the same way as in bona fide Tilman's diagram, the species $B_\alpha$ can coexist with the species $B_\beta$ if the one-microbe line of $B_\alpha$ ends at the point at which $B_\beta$ can grow, and vice versa (see Fig. S13, panel A). If however the microbe's one-microbe line lands on the ZNGI at a point where for any other competitor the growth is impossible, it becomes the sole winner (barring the bistability cases), as it is shown on the panels B, C Fig. S13. Note that the one-microbe line of $B_\beta$ always lands on the microbe's ZNGI at larger values of $T_1$, as its growth rate $g_{\beta 1}$ is smaller, and thus its $T_1 = (1/g_{\beta 1}) \log \Big( (D R_1 + R_2)/(R_1 + R_2) \Big)$ is larger than the depletion time of $R_1$ from (S.26). This is why the one-microbe line of $B_\beta$ is always shown on the right of that of $B_\alpha$. (Remember that we consider yields of different microbes wrt different sources to be equal. If they are not equal, the one-microbe lines will display more complex behavior, see the corresponding SI section.)

Another thing worth mentioning is that $T_1$ in Eq. (S.26) obviously grows larger with the increase of the ratio $R_1/R_2$, thus moving the steady state point down the slope (or up the slope, if the microbe is anomalous) of the corresponding ZNGI in the Time plane.

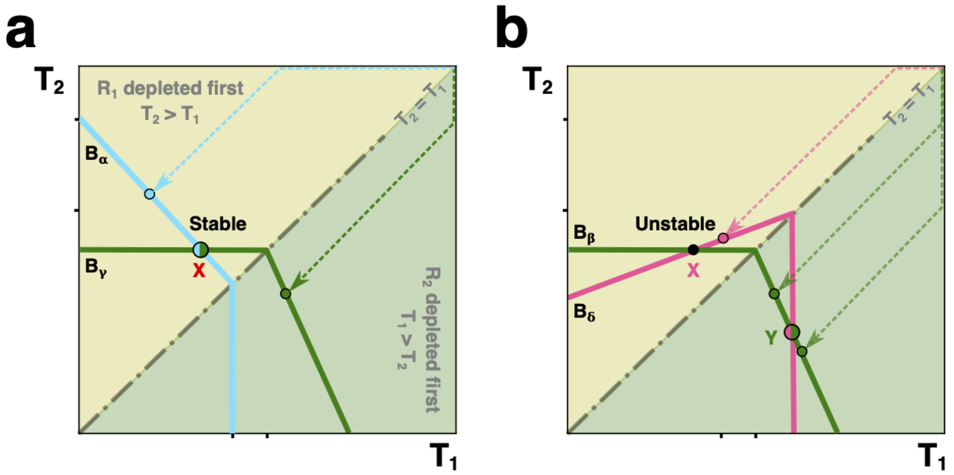

Figure S14: **Stable (a) and unstable (b) coexistence. (a)** The (light blue, dotted) one-microbe line of the bacterium $B_\alpha$ lands on its (light blue, solid) ZNGI at the point where $B_\gamma$ can grow. The (green, dotted) one-microbe line of the microbe $B_\gamma$ lands on its (green, solid) ZNGI at the point where $B_\alpha$ is able to grow. The invasion of the environment by both microbes will result in stable coexistence. **(b)** The (pink, solid) ZNGI of the anomalous microbe $B_\delta$ (top choice $R_1$) intersects with the (green, solid) ZNGI of the normal microbe $B_\beta$ (top choice $R_2$) at points **X** above the diagonal $T_1 = T_2$ and **Y** below the diagonal. To achieve coexistence, the one-microbe lines of each microbe should land where the other one can grow. The one-microbe line of the anomalous $B_\delta$ (pink, dotted) lands on the microbe's ZNGI (pink, solid) at a point where $B_\beta$ can grow. If the (green, dotted) one-microbe line of $B_\beta$ lands on its (green, solid) ZNGI above the point **Y**, where the anomalous $B_\delta$ cannot grow, then the normal $B_\beta$ comes out a sole winner. If the one-microbe line of $B_\beta$ lands on its ZNGI below the point **Y**, where $B_\delta$ can grow, both species would end up coexisting at the point **Y**. The coexistence at the point **X** thus cannot be realizable by competition, and is unstable if prepared.

#### B.4 Types of the steady state coexistence: stable and unstable

The coexistence shown in Fig. S13 is obviously stable as no single one of the given two microbes competing for the resources cannot on its own supplant another. The same goes for the coexistence of the species $B_\alpha$ and $B_\gamma$ that have different preferences with regard to the resources $R_1$ and $R_2$, see Fig. S14, panel A. Coexistence at the point $X$ is stable when the one microbe line of $B_\alpha$ hits it ZNGI above the point $X$ (and thus the zero net growth isocline of its rival $B_\gamma$ goes under that of $B_\alpha$) and the same is true for the one microbe line of $B_\gamma$, so that at the point of its own single-microbe steady state it cannot supplant $B_\alpha$. This (panel A) is the case when both microbes are normal. (One is reminded that we consider a microbe "normal" if it grows fastest on its top choice. Otherwise, the species is regarded as"anomalous".) On the contrary, when the anomalous species enter the picture (Fig. S14, panel B), some steady states that one can technically prepare turn out to be unstable. For instance, the coexistence of the microbes $B_\gamma$ (normal) and $B_\delta$ (anomalous) at the point $X$ (Fig. S14, panel B) would not work; if prepared, it will degenerate either into the coexistence point $Y$, if the one-microbe line of $B_\gamma$ goes the way the solid version (1) does, landing on its own ZNGI where $B_\delta$ can grow, or else into the single-microbe steady state of $B_\gamma$ if the latter's one-microbe line goes like the dashed version (2). (If the one-microbe line of $B_\delta$ lands to the left of the point $X$, then $B_\delta$ is the sole winner.) That stands to reason from the physical point of view. Prepare the steady state at the point $X$ and disturb the environment, say, by adding a small amount of the species $B_\gamma$ making its abundance a little larger than that required for the steady state. That would not affect at first the depletion time $T_1$ of $R_1$ as $B_\gamma$ does not consume it in the state in question, yet the depletion time $T_2$ in the dilution cycle during which the abundances got disturbed will be shorter, as there is more of $B_\gamma$ to use it up. As a consequence, $B_\delta$ will come out of it grown less than $D-$fold, and for the next cycle its abundance will be less than the

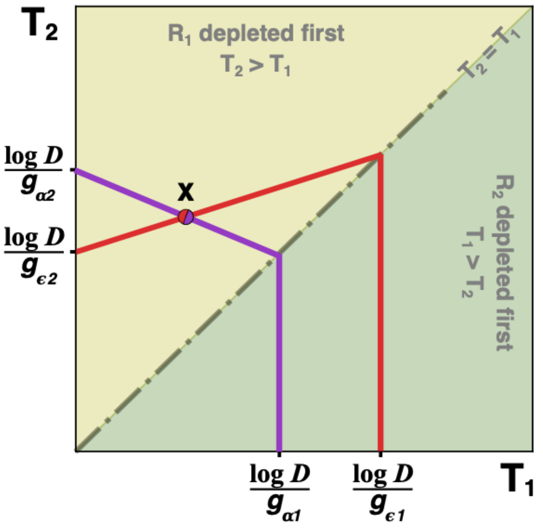

Figure S15: **An anomalous species coexisting with a normal one with the same nutrition preferences.** Within a particular range of the values of the ratio $R_1/R_2$ the anomalous bacterial species $B_\epsilon$ (the red ZNGI with the "anomalously" positive slope above the diagonal $T_1 = T_2$) can coexist with the normal microbe $B_\alpha$ (the lilac ZNGI with the "normal" negative slope above the diagonal) at the point **X**.

steady state amount. So it will be slower yet to consume $R_1$ at which it grows less fast than at $R_2$ being anomalous, and $B_\gamma$ will have more time to grow on its favored resource $R_2$ all on its own. Thus $B_\delta$ gets lesser share of $R_2$ that is essential for its growth and loses the next dilution cycle, and so on. Eventually the depletion order changes to first $R_2$, then $R_1$, and the outcome depends on whether $B_\gamma$ can supplant $B_\delta$ (dashed line) or will have to share (solid line in Fig. S14, panel B). This is determined by the ratio $R_1/R_2$ (see e. g. eq. S.25 or the discussion in previous sections).

#### C Anomalous microbes and their rarity

In this section we discuss the anomalous species. Generally speaking, if a bacterium $B_\epsilon$ with a diauxie growth curve prefers the resource $R_i$ to the resource $R_j$ despite the fact that it grows faster on the less-preferred nutrient, $g_{\epsilon i} < g_{\epsilon j}$, we consider it anomalous.

The definition would stay rather vague unless we claim the absolute knowledge of the preference list of a species in its totality. Effectively, we would deal with a limited number of resources and will probably not notice any effect of a species being anomalous unless it is its top choice (among the nutrients available) that provides the microbe with a lesser value of the growth rate than the second-best. This is the context in which the term "anomalous" will be used further on, the more so as, in this section, we deal mostly with an artificial environment containing just two resources.

In our simulations, using the balanced nutrient supply (meaning that the concentrations of the resources are equal at the beginning of each dilution cycle) and the values of the dilution coefficient $D$ varying from 2 to 10000, we find the anomalous microbes vanishing from the steady state community of survivors as long as the latter becomes more and more mature.

To see why that happens, consider point $X$ in Fig. S15, which is a point in the Time plane at which the microbe $B_\alpha$ that is normal and prefers $R_1$ to $R_2$ coexists with the microbe $B_\epsilon$ which is anomalous, although it shares the former's preferences. If in the steady state $T_1 < T_1(X)$, the anomalous species is the winner, and we are interested in the values of the ratio $R_1/R_2$ and the dilution coefficient $D$ at which that happens. Note that $g_{\epsilon 1} < g_{\alpha_1}$ meaning that the anomalous microbe we consider here does not excel at both growth parameters, only at that of its second choice. Thus our case has nothing to do with the "superbug" event, we consider a regular situation.

Consider $\Delta T_1 = T_1 - 0$ and $\Delta T_2 = T_2 - T_1$, the actual lifetimes of the resources $R_1$ and $R_2$ from the moment they, each in its turn, are attacked by the consuming microbe(s). If $B_\epsilon$ is a single tenant that had established the steady state and $B_\alpha$ is invading, the latter cannot survive when

$$g_{\epsilon 1}\,\Delta T_1 + g_{\epsilon 2}\,\Delta T_2 > g_{\alpha 1}\,\Delta T_1 + g_{\alpha 2}\,\Delta T_2. \tag{S.27}$$

$\exp(g_{\epsilon 1}\Delta T_1) = R_1 + N_\epsilon$ for the moment when $R_1$ is totally consumed, $N_\epsilon$ being the steady state concentration of the bacterium $B_\epsilon$ at the beginning of each dilution cycle, and the steady state multiplication rule $g_{\epsilon 1}\Delta T_1 + g_{\epsilon 2}\Delta T_2 = D$, provide us with the values of the effective times of consummation for both resources:

$$\begin{cases} \Delta T_1 = \dfrac{1}{g_{\epsilon 1}} \log\left(\dfrac{DR_1 + R_2}{R_1 + R_2}\right) \\[2ex] \Delta T_2 = \dfrac{1}{g_{\epsilon 2}} \log\left(\dfrac{D\,(R_1 + R_2)}{DR_1 + R_2}\right) \end{cases} \tag{S.28}$$

Combining (S.27) and (S.28) we get

$$\frac{g_{\alpha 1} - g_{\epsilon 1}}{g_{\epsilon 1}} \log\left(\frac{DR_1 + R_2}{R_1 + R_2}\right) < \frac{g_{\epsilon 2} - g_{\alpha 2}}{g_{\epsilon 2}} \log\left(\frac{D\,(R_1 + R_2)}{DR_1 + R_2}\right). \tag{S.29}$$

As long as (S.29) holds, the normal microbe $B_\alpha$ cannot invade the environment shaped by the anomalous microbe $B_\epsilon$. To get the criterion for the event of $B_\epsilon$ withstanding the invasion in a convenient form, we transform (S.29) and introduce a parameter

$$y = \left(\frac{g_{\epsilon 2}(g_{\alpha 1} - g_{\epsilon 1})}{g_{\epsilon 1}(g_{\epsilon 2} - g_{\alpha 2})} + 1\right)^{-1} = \frac{g_{\epsilon 1}\,(g_{\epsilon_2} - g_{\alpha_2})}{g_{\epsilon 2} g_{\alpha 1} - g_{\epsilon_1} g_{\alpha 2}} < 1. \tag{S.30}$$

We get the criterion

$$\frac{R_1}{R_2} < \frac{D^y - 1}{D - D^y} \tag{S.31}$$

The criterion (S.31) can mean either a huge imbalance in terms of the nutrient supply or something reasonable, depending on the values of $y$ and $D$.

Using elementary calculus, we conclude that the right hand part of the equation is a decreasing function of $D$, at least when it goes all the way from 2 to $\infty$, at any fixed $y$.

While at $D = 2$ it is possible to get the left-hand part of (S.31) well over $R_1/R_2 = 1$ at reasonable values of $y$, there is yet another factor coming into the game, closely related to the rise of complementarity at any value of $D$. Consider the diagram from Fig. S15 with the addition of yet another species $B_\gamma$ that prefers $R_2$ to $R_1$ and is a capable consumer of its top choice.

In Fig. S16 the green species $B_\gamma$ invades the environment shaped by the coexisting $B_\alpha$ the normal and $B_\epsilon$ the anomalous microbes. The invader is perfectly able to grow as its lifeline passes below $X$ that is the point of the said coexistence. When $B_\epsilon$ grows, it affects $T_2$ that is becoming less, meaning that the microbes $B_\alpha$ and $B_\epsilon$ start each dilution cycle in smaller amounts and $T_1$ becomes larger. That shifts the position of the system towards the point $Z$ of the coexistence of two normals $B_\alpha$ and $B_\gamma$ that have different nutrition preferences.

One might wonder whether the complementarity manifests itself at all at the smaller values of the dilution coefficient $D$. The question is closely related to the crucial importance of the top choice nutrient and the value of the growth rate of the microbe associated with it. One of our arguments for the top choice being the most important from the very beginning was that in order to reach the steady state, a species must have its initial concentration multiplied by $D$

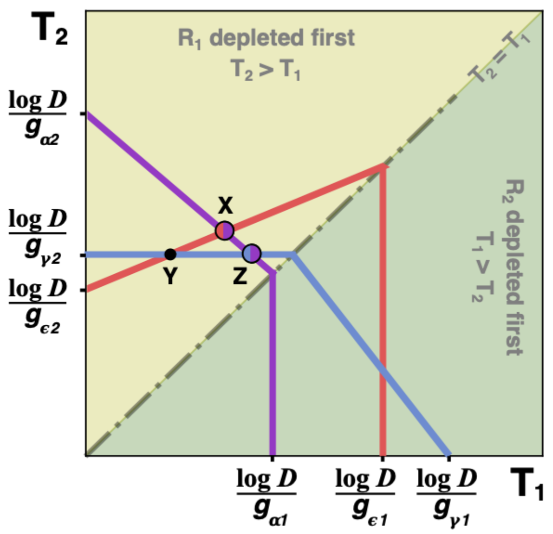

Figure S16: **The coexistence of the anomalous species $B_\epsilon$ with the normal species $B_\alpha$ with the same preferences disabled by an invasion of the microbe $B_\gamma$ with the reversed preferences and thus giving way to the complementarity.** An anomalous microbe $B_\epsilon$ (red ZNGI with the positive slope above the diagonal $T_1 = T_2$, top choice $R_1$) can coexist with a normal microbe $B_\alpha$ (lilac ZNGI, top choice $R_1$) at the point **X** in a stable manner. However, the microbe $B_\gamma$ (light blue ZNGI, top choice $R_2$) with the complementary food preferences can invade at that point and supplant the anomalous microbe, switching the state of the system to the point **Z** at which the normal microbes $B_\alpha$ and $B_\gamma$ share the environment. The coexistence point **Y** of $B_\gamma$ and $B_\epsilon$ is unstable.

towards the end of each dilution cycle, and most of that multiplication is done at the expense of the first nutrient it consumes. Indeed, the steady state value of the microbe concentration when it is the single tenant of the environment being $\sum R_i/(D-1)$, which is also $nR/(D-1)$ when all the $n$ resources are present in equal concentrations, is much less than $R$ if $n \ll D$. So, upon consuming its top choice, a nutrient has its concentration multiplied by $(D-1)/n$ which is still a huge number, while the second nutrient in its turn multiplies it by less than 2, and each of the next-in-line resources does even poorer job at multiplying. On the contrary, at $D = 2$, the first resource augments the initial concentration of the microbe by a fraction $1/n$. While the second nutrient provides even less than that in terms of multiplication, that particular argument in favor of the top choice being the most important one becomes much less convincing.

What matters for the comparative importance of the first, second and further choices, as well as for the rise (or fall?) of complementarity, is the fact that the nutrient which is the last to be depleted remains the most reasonable target for invasion. Moreover, in the case of maximum diversity of the steady state community, complementarity is either already present or can be produced by an invasion with the probability that does not depend on $D$ but rather is determined by the properties of the particular growth rate distributions. In the case of two nutrients, the equilibrium differential times are determined by the system of equations:

$$\begin{cases} g_{\alpha 1}\,\Delta T_1 + g_{\alpha 2}\,\Delta T_2 = \log D \\ g_{\beta 1}\,\Delta T_1 + g_{\beta 2}\,\Delta T_2 = \log D \end{cases} \tag{S.32}$$

(which in the general case would contain $n$ equations for $n$ unknowns), the solution would obviously not depend on the nutrient concentrations, and the condition for an invader $B_\gamma$ whose top choice is $R_2$ being able to successfully invade would have the form:

$$g_{\gamma 2} > \frac{g_{\alpha 1}\,g_{\beta 2} - g_{\alpha 2}\,g_{\beta 1}}{g_{\alpha 1} - g_{\alpha 2} + g_{\beta 2} - g_{\beta 1}} \tag{S.33}$$

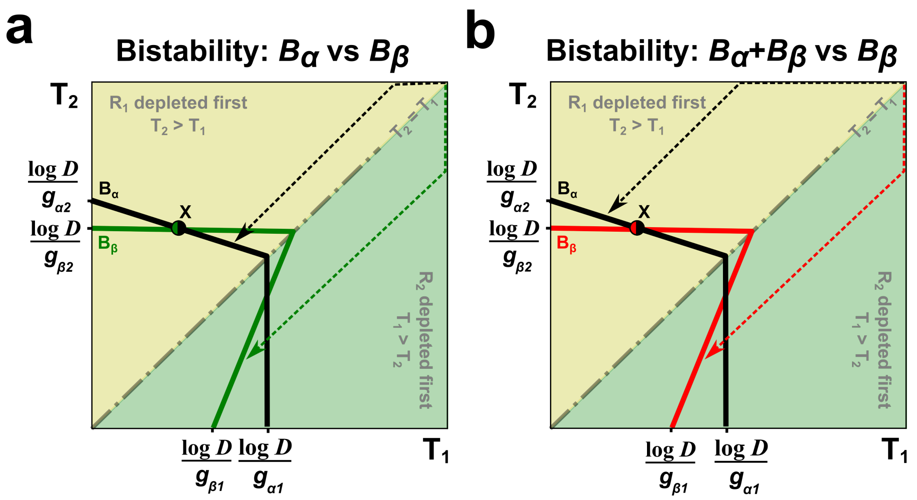

Figure S17: **Bistabilities.** **(a)** The (black, dotted) one-microbe line of $B_\alpha$ lands on the microbe's (black, solid) ZNGI at the point where the bacterium $B_\beta$ cannot grow. The (green, dotted) one-microbe line of $B_\beta$ land on its (green, solid) ZNGI where the species $B_\alpha$ cannot grow. The resulting steady state will depend upon which microbe was the first to invade, hence there is bistability between two possible outcomes with different depletion orders. **(b)** The (dotted, red) one-microbe line of $B_\beta$ lands on the microbe's (solid, red) ZNGI at the point where $B_\alpha$ cannot grow. The (dotted, black) one-microbe line of $B_\alpha$ lands on its (solid, black) ZNGI at the point where $B_\beta$ can grow on its top choice, never displaying a diauxie shift. The resulting steady state will depend upon which microbe was the first to invade. If the bacterium $B_\alpha$ comes first, then, upon the invasion of $B_\beta$, the system acquires the steady state with both bacteria coexisting at the point **X**. If the microbe $B_\beta$ is the first to invade, it will remain the sole resident. Hence there is bistability between two possible outcomes with different depletion orders and different degrees of diversity.

It can be easily seen that the invader having its $g_{\gamma 2}$ equal (or less than but close) to $g_{\alpha 1}$ is the definite winner. Substituting the index $\beta$ with $\epsilon$ we get the condition under which the invader $B_\gamma$ chases away the anomalous species $B_\epsilon$. The anomaly of $B_\epsilon$, if anything, further frees the value of $g_{\gamma 2}$. (Of course, when $g_{\gamma 2} > g_{\alpha_1}$, the invader wins hands down, but in this case the depletion order will get reversed, and we would have to consider the anomalous species with the same preferences as $B_\gamma$ all over again.)

To summarize, the anomalous species get driven to extinction at any values of the dilution coefficient $D$ barring the cases of a huge imbalance in the nutrient supply, and the complementarity in a mature steady state also persists regardless of whether $D$ is 2 or 10000.

### D Multistability in a community of diauxic shifters

#### D.1 What prompts multistability

In some cases more than one uninvadable steady state is possible for the same initial conditions. Depending on the order of invasions, the system in the Time plane (or Time space, when the sources of nutrition are multiple) might end up at different points of the Pareto-optimal boundary. In the models associated with the continuously diluted chemostat, the phenomenon of multistability has been known to occur[5, 6]. With the co-utilisers, it comes to light when different yields of different microbes using the same resource are taken into account. When one deals with the diauxic shifters, however, the conflict of the preference lists of microbes and the rank lists of their growth rates, in other words, the presence of the anomalous microbes, is enough to produce multistability.

#### D.2 An example of bistability between two single-microbe states

In the case when only two microbes are present in the environment, bistability can occur. The reason why multistability can arise is easily seen by considering that simplest example.

In Fig. S17 (a) the black arrow points at the place at which the single-microbe state I of the (normal) microbe $B_\alpha$ is located on its zero net growth isocline (also black), and the green arrow marks the place at which the single-microbe state II of the (anomalous) microbe $B_\beta$ hits its own (green) ZNGI. When $B_\alpha$ has shaped the environment in its single-microbe steady state I, $B_\beta$ cannot invade, and vice versa.

To get the condition under which that kind of bistability can be realized, write for the environment shaped by $B_\alpha$ (with the depletion times $T_1^{(\alpha)} < T_2^{(\alpha)}$, $T_1^{(\alpha)} = (1/g_{\alpha 1}) \log((DR_1 + R_2)/(R_1 + R_2))$, $T_2^{(\alpha)} = T_1^{(\alpha)} + (1/g_{\alpha 2}) \log(D(R_1 + R_2)/(DR_1 + R_2)))$

$$g_{\beta 2} T_2^{(\alpha)} < \log D, \tag{S.34}$$

and for the environment shaped by $B_\beta$

$$g_{\alpha 1} T_1^{(\beta)} < \log D, \tag{S.35}$$

with the depletion times $T_1^{(\beta)} > T_2^{(\beta)}$, $T_2^{(\beta)} = (1/g_{\beta 2}) \log((DR_2 + R_1)/(R_1 + R_2))$, $T_1^{(\beta)} = T_2^{(\beta)} + (1/g_{\beta 1}) \log(D(R_1 + R_2)/(DR_2 + R_1))$.

From (S.34, S.35) we get the bistability condition:

$$\frac{R_1}{R_2} > \max\left(\frac{D^x - 1}{D - D^x}, \frac{D - D^y}{D^y - 1}\right), \tag{S.36}$$

where

$$x = \frac{g_{\alpha 1}}{g_{\beta 2}} \frac{g_{\beta 2} - g_{\alpha 2}}{g_{\alpha 1} - g_{\alpha 2}}, \tag{S.37}$$

and

$$y = \frac{g_{\beta 2}}{g_{\alpha 1}} \frac{g_{\beta 1} - g_{\alpha 1}}{g_{\beta 1} - g_{\alpha 2}}. \tag{S.38}$$

##### D.3 An example of bistability between coexistence and exclusion for two microbial species

If the anomalous species $B_\beta$ is very good at consuming both nutrients, the bistability may occur between the state at which both $B_\alpha$ and $B_\beta$ are present peacefully coexisting — and the state at which it's $B_\beta$ alone that has won the game.

Such a situation is presented in Fig. S17 (b). Here, the condition under which that bistability can be realized is more tricky

$$\frac{D^x - 1}{D - D^x} > \frac{R_1}{R_2} > \frac{D - D^y}{D^y - 1}, \tag{S.39}$$

with $x$, $y$ the same as above. That implies $(D^x - 1)/(D - D^x) > (D - D^y)/(D^y - 1)$, i. e. $D^x + D^y > D + 1$. At least one of $x$, $y$ would have to be rather close to 1, to make that happen.

##### D.4 Conclusions

In the line of thought of a recent work[7] that names conflicts as the source of complexity, in our system multiple steady states at the given external conditions occur when there is a conflict between the preference order inherent to a microbe and the rank order on its growth rate. We do not see much of that in our simulations of the environment with the balanced nutrient supply, although with an anomalous "super-bug" in the pool that could be arranged.

### E On the attainability of the final steady state: does the invasion process end?

There is yet another complication to our model of the serial invasion related to the anomalous species. Why should the final steady state, uninvadable by any microbe from a given pool, even exist? We could have famous "rock-paper-scissor" cycles or stochasticity, even though we have never observed them in our simulations. In this section, we address two questions: (1) whether the situations in which the invasion process never ends can be arranged within the frames of our model; (2) if the answer is positive, why then do we always get a final and uninvadable state in our simulations.

#### E.1 An example of a cyclic final solutions

When points of unstable coexistence like $X$ in Fig. S14 (b) appear on the Pareto-optimal boundary, they become "percolation points" of sorts. The dynamic trajectory of the system might "flow" from them over the line $T_1 = T_2$. Indeed, when $B_\gamma$ invades the environment shaped by $B_\alpha$ (Fig. S14 (a)), it gets arrested at the point $X$, and the depletion order remains first $R_1$, then $R_2$, while invading the environment shaped by $B_\delta$ (Fig. S14 (b)) it either cannot survive or, never getting stuck at the point $X$, brings the system to some point where $T_1 > T_2$. That is almost of no consequence in the case of the environment containing just two resources, but when there are more of those, all the relevant points of coexistence, technically, can be unstable. When the latter is the case, we might see cycles or even stochasticity instead of the final uninvadable steady state.

An example of a cycle of the rock-paper-scissors type can be given for an environment containing three sources $R_1, R_2, R_3$, say, in equal abundances $R_1 = R_2 = R_3 = R$. Take three anomalous species $B_\phi, B_\psi, B_\chi$ differing in their preferences only, so that $g_{\phi 1} = g_{\psi 2} = g_{\chi 3} = xg$, $g_{\phi 2} = g_{\psi 3} = g_{\chi 1} = g$, $g_{\phi 3} = g_{\psi 1} = g_{\chi 2} = yg$, $x, y < 1$. (For $B_\phi$, the preferential order goes $R_1, R_2, R_3$, in the case of $B_\psi$ it's $R_2, R_3, R_1$, and $B_\chi$ would have it as $R_3, R_1, R_2$.) The microbes thus have the same set of growth rates: $xg$ on their first choice, $g$ (the maximum) on the second and $yg$ on their third. Introduce for the sake of convenience

$$\tau_1 = \frac{1}{g} \log \frac{D+2}{3}, \; \tau_2 = \frac{1}{g} \log \frac{2D+1}{D+2}, \; \tau_3 = \frac{1}{g} \log \frac{3D}{2D+1}, \tag{S.40}$$

so that in an environment shaped by, say, $B_\phi$ the depletion times will be

$$\Delta T_1 = T_1 = \frac{\tau_1}{x}, \; \Delta T_2 = T_2 - T_1 = \tau_2, \; \Delta T_3 = T_3 - T_2 = \frac{\tau_3}{y}. \tag{S.41}$$

Any of the other two anomalous competitors would, on its own, shape the environment in a way identical to the above up to cyclic permutation. Find the values of $x, y < 1$ such that $B_\psi$ (the consummation order $R_2, R_3, R_1$) can supplant $B_\phi$ (c. o. $R_1, R_2, R_3$) upon invasion while $B_\chi$ (c. o. $R_3, R_1, R_2$) would not be able to invade. $B_\psi$ being able to invade means $g_{\psi 2} T_2 + g_{\psi 3}(T_3 - T_2) > \log D$, i. e.

$$xg\,(\frac{\tau_1}{x} + \tau_2) + g\,\frac{\tau_3}{y} > \log D = g\,(\tau_1 + \tau_2 + \tau_3) \tag{S.42}$$

From the above we get

$$\frac{1-y}{y\,(1-x)} > \frac{\tau_2}{\tau_3} \simeq 1.7 \tag{S.43}$$

(the last evaluation we get assuming $D \gg 1$, so that $g\,\tau_2 \simeq \log 2$, $g\,\tau_3 \simeq \log 1.5$ as it is evident from (S.40)). Now, to enable the rock-paper-scissors process, one needs to ensure that $B_\chi$ cannot invade the environment shaped by $B_\phi$ (while $B_\psi$ can if the condition (S.43) holds, and does chase $B_\phi$ out upon invasion).

$$g_{\chi 3}\, T_3 = x\, g\, (\frac{\tau_1}{x} + \tau_2 + \frac{\tau_3}{y}) < \log\, D = g\, (\tau_1 + \tau_2 + \tau_3), \tag{S.44}$$

meaning

$$\frac{x - y}{y\, (1 - x)} < \frac{\tau_2}{\tau_3} \simeq 1.7 \tag{S.45}$$

Combining (S.43) with (S.45) we get the rock-paper-scissor condition for the environment that gets repeatedly invaded by the three species $B_\phi, B_\psi, B_\chi$:

$$\frac{x - y}{y\, (1 - x)} < \frac{\tau_2}{\tau_3} \simeq 1.7 < \frac{1 - y}{y\, (1 - x)} \tag{S.46}$$

The above gets easily satisfied. Indeed, take $x = 0.7$, $y = 0.5$ and get $4/3 < 1.7 < 10/3$ which is obviously true. This means we have organized a rock-paper-scissor cycle: whenever $B_\phi$ installs itself as the resident, $B_\chi$ trying to invade will not succeed, but $B_\psi$ will. However, having chased out $B_\phi$, the winner $B_\psi$ will shape the environment so that, due to the symmetry of the system parameters, it will eventually get chased out in its turn by $B_\chi$, and the latter, upon establishing its residence, will get chased by the $B_\phi$; at that point, the cycle gets completed and the story will go on.

The question whether a cycle can be organized by the "pedantically normal" species with the preferential list mirroring the table of ranks of their own growth rates, i. e. those that always prefer a nutrient $R_i$ to the nutrient $R_j$ if they grow faster on $R_i$, we leave unanswered. Though it is interesting in itself and the answer is probably no, to all the real as well is to our modeling purposes the question is academic. We consider a species normal if its top choice is the nutrient on which it grows faster than on all the other nutrients present in the environment. The definition of normality / anomaly is thus environment-dependent: a microbe that seems perfectly normal when its absolute top-choice is present may behave weirdly when the latter is absent (or vice versa). We are perfectly fine with this dependence as it helps us single out a way in which the evolution pressure might go in a particular environment. When we use our definition, the example above is easily modified to present a case in which normal microbes can form a cycle. Add to the environment a universally loved nutrient like glucose and ascribe to all the microbes from the example above the same growth rate $G > g$ on it, making it their first choice. The consideration goes then very similar to the above, although the numerical criterion changes, but the possibility of the cycle remains.

We do not see cycles (or stochasticity, for that matter) in our simulations because the anomalous species vanish from the community as the latter matures, and thus the Pareto-optimal boundary contains only stable steady state points. For a more detailed explanation see below.

#### E.2 Why the invasion process ends

Suppose there is an endless cycle or stochastic sequence of the steady states (supplanting each other as a result of a successful invasion) that cannot be broken down to an uninvadable state by a series of out-of-turn invasions. By the right of random chance, we are allowed to optimize it: if a steady state 1 is encountered more than once and the states that replace it in the given endless sequence are multiple: 2, 3, ..., then upon encountering the state 1 we always choose the unique invasion that results in the least overall depletion time of the state replacing 1 at this point. As the number of the steady states is finite, and in the end of the optimization each state is supplanted by a particular one, we turned the sequence into a cycle. It is also safe to assume that all the anomalous species are already banned from the community of survivors.

Our argument is, basically, the following. In the mature community all the depletion times are close to $\log D/G$, where $G$ is a typical growth rate of an expert on its top choice. (Here we call a species "expert" or "champion" of its favorite resource $R_i$ if it is (a) normal and (b) best-growing on $R_i$ among all the normals that share that same top choice.) The cycle is bound to include some invasion events in which a new resident chases out some of the old ones. Usually the successful invader has a higher growth rate on its top choice than the one that is chased out. That however cannot be always the case as it is impossible to have a cycle with the net growth rate on the top choice increasing at each step. So at some point $B_\psi$ with the top choice $R_i$ supplants $B_\phi$ that prefers $R_j$ even though $g_{\phi j} < g_{\psi i}$ (here some caveats are needed, and we'll get to them later). A crude approximation would have $T_i \sim T_j \sim T \sim \log D/G$, where $T$ is the overall effective growth time; all differences between depletion times $\sim (\Delta G/G^2) \log D$ (where $\Delta G = G_{max} - G_{min}$, $G_{max}$ being the growth rate of the absolute fastest growing on its own top choice champion (or expert), $G_{min}$ — the growth rate of the slowest one among the champions). We would also introduce $\langle g_\phi \rangle$ and $\langle g_\psi \rangle$ as average, in a sense, growth rates of $B_\phi$ and $B_\psi$ on their second choices and further on, so that we describe the invasion event as

$$g_{\phi j} T_j + \langle g_\phi \rangle (T - T_j) > g_{\psi i} T_i + \langle g_\psi \rangle (T - T_i). \tag{S.47}$$

Resorting to the crude approximations of the differences between times and growth rates, we transform the above into

$$(\langle g_\phi \rangle - \langle g_\psi \rangle) \frac{\Delta G}{G^2} \log D > \Delta G \frac{\log D}{G} \tag{S.48}$$

meaning

$$\langle g_\phi \rangle - \langle g_\psi \rangle > G, \tag{S.49}$$

something that is not likely to happen unless the species $B_\phi$ is anomalous. There are of course those caveats, one of them being that the overall depletion time, though small, can still happen to be larger than $\log D/G_{min}$ (though the worst that can actually happen to it is being of the order $\log D/G + \log(n/(n-1))/\langle g \rangle$ if the last nutrient to be depleted is not occupied, as all the microbe residents having feasted on their favorite choices join in consuming the last unoccupied nutrient with their total concentration over $(n-1)R$, growing upon it with some average rate $\langle g \rangle$, so that $(n-1)R \exp(\langle g \rangle \Delta T_{last}) \sim nR,$), $n$ being the number of resources present in the environment and $\Delta T_{last}$ – the short length of time through which only the last nutrient alone is left in the environment. The other is that the invasion event at which the average residential top-choice growth rate gets diminished can be the invasion of the champion of the last nutrient to be depleted with no resident chased out. Still, if that happens, after that one would have to have yet another invasion diminishing the average top-choice growth rate of the first (n - 1) resources, as that one can not grow forever, and the inequality of the same sort as (S.47) will be valid. Even if the invader $B_\psi$ should actually be very good (on the expert level) at growing on its first two nutrients and not necessarily anomalous, it is still a lot to ask of a chance. By increasing the number of species, the probability to get a top choice on the expert level is made smaller and the probability to bring the consummation of two first choices to the expert level plummets down as a square of the former.

All the above is true of course only for the case of a reasonably balanced nutrient supply, while huge imbalance in the resources initial abundances will be favorable for the anomalous species that are good enough and start with the resource which is scarce and excel in their second choice which is abundant (in this way, they are certain to grow mostly on their second choice). The cycle of the sort discussed in the previous section will be hard to organize in this case as it would need certain symmetry, but some complicated way over the Pareto-optimal boundary through its "percolation points" (unstable coexistence points) may be possible.

Even a balanced environment may have a history of this or that resource getting scarce, and for a while can be occupied by the family of the anomalous species. Within such a family, stochasticity can arise, and increase the effective diversity to the point of breaking the rule of the number of the survivors being less than or equal to the number of resources. After all, that rule is valid for the steady states and not for the effective collection of those.

### F On the number of surviving species

#### F.1 Steady state survivors

The usual rule (related to the competitive exclusion principle) stating that "the number of surviving species is less or equal to the number of resources" is applicable to the steady states in the serial dilution experiments, whenever the external conditions are fixed. As always, varying the resource concentrations and / or the dilution coefficient we move the system over the Pareto-optimal boundary, so that different uninvadable steady state communities may assemble themselves under different external conditions.

#### F.2 Why the competitive exclusion principle applies

Any steady state is associated with a particular depletion order. When only two nutrients are present in the environment, the Time Plane $(T_1, T_2)$ is divided into two regions: above the diagonal $T_1 = T_2$, with $T_1 < T_2$ and consequently the resource $R_1$ being depleted first, and below the diagonal, with $T_1 > T_2$ hence the reversed depletion order. In the same way, when there are $n > 2$ resources and one has to consider the $n-$dimensional Time Space instead of the Time Plane, the relevant (with all the time coordinates positive) part of the former gets divided by the hyperplanes $T_i = T_j$ into $n!$ segments, each corresponding to a particular depletion order. As long as the depletion order is decided upon, the condition of survival for any microbe can be written in the form of a linear equation. Suppose, for instance, that the depletion order is $T_1, T_2, ..., T_n$ and the bacterium $B_\alpha$ has the preference list beginning with $T_1, T_4, T_{n-2}, T_n, ...$. In this case, in the particular segment of the Time Space the condition for the microbe's survival will be:

$$g_{\alpha 1}\, T_1 + g_{\alpha 4}\, (T_4 - T_1) + g_{\alpha, n-2}\, (T_{n-2} - T_4) + g_{\alpha n}\, (T_n - T_{n-2}) = \log\, D$$

We get the number of the survival conditions equal to the number of competing species from the pool $N \gg n$, while the number of unknown depletion time variables is equal to $n$. Thus obviously, in the general case, no more than $n$ equations can hold simultaneously. There are additional conditions like positiveness of all solutions, uninvadability of a solution, the question of whether the obtained solution is adequate, meaning that it is located on the real and not on the virtual segment of the Pareto-optimal boundary. (To wit, if we get, say, a "single-microbe steady state" corresponding to $B_\omega$ growing all the time on its preferred nutrient $R_n$ because it happened to be depleted last, we know that cannot be a real-life solution, as, if left alone, this microbe will eventually change the depletion order by shaping the environment in accordance with its own preference list.) In any case, only a degeneration in the set of the growth rate parameters can lead to surviving of more than $n$ species at the same time ($n$ being the number of the resources present in the environment).

#### F.3 Trade-off miraculous diversity boost takes too much to be manifested

One specific case of such a degeneration is that of the "trade-off" well known to the researchers familiar with the MacArthur model. If, so to speak, a species can increase its growth rate on a certain source only by decreasing its growth rate on some other nutrients, so that all the microbes in the pool have the same average growth rate, there is a possibility that an arbitrary number of species can coexist, if the conditions are right. Namely, if, for example, the depletion order is $R_1, R_2, ..., R_n$ and the depletion times go as $T_1 = T_2 - T_1 = T_3 - T_2 = ... = T_n - T_{n_1} = \Delta T$, all the microbes with the growth rates constrained by the trade-off condition *and having the same preferential order* in principle can coexist. If the environment is shaped by $B_\alpha$ so that

$$g_{\alpha 1}\, T_1 + g_{\alpha 2}\, (T_2 - T_1) + ... + g_{\alpha n}\, (T_n - T_{n-1}) = (g_{\alpha 1} + g_{\alpha 2} + ... + g_{\alpha n})\, \Delta T = \log\, D,$$

the same equality will hold for any of the other bacteria sharing that same preferential order and subject to the same trade-off in their growth rates. When added in a seed amount, an invader, though not really being able to grow, will not die out. Although such a steady state can be theoretically prepared (for any, arbitrarily chosen in advance, set of initial concentrations of the participating microbes), it would be an exotic situation: first, $R_2$ should be much more abundant than $R_1$ and $R_3$ much more abundant than $R_2$ and so on, the abundances of the sequent nutrients would have to form something close to a geometric progression. Second, the microbes participating in such a community should have strictly the same preferential order; an invader with the same average growth rate but with different preferences would easily win the game as the trade-off will not present to it much of a constraint. This, by the way, should be the case of concern for the MacArthur model as well: if you have the trade-off averaged over, say, 10 resources and there are only three of them in the environment, then the trade-off will not have the chance to manifest itself and the results you get would look as if there was not any.

##### F.4 Conclusion: the regular rules of the competitive exclusion apply

To conclude, in this paper we deal with a regular computer-resource model for which the rule of "the number of surviving species being less or equal to the number of resources" is applicable as long as it is a steady state and not some cycle or stochastic process at the end of the system's maturation.

#### G How different yields affect the outcome of the serial dilution experiment

Throughout the main text we assumed all the yields of all the bacterial species wrt different resources to be equal. This however doesn't have to be the case. The effects of differences in terms of yields between species (and, for a single species, wrt different resources) have been known to include multistability [4] and other indications of growing complexity, like "rock, paper, scissors" cycles or stochasticity [8] in the system. In our model, even assuming all the yields equal, we get the above effects when the anomalous microbes manifest themselves. Yet, as we have shown, in the regular situations the anomalous species tend to die out. Accounting for the difference in yields however brings multistability back in the picture.

Dynamics of the environment containing one resource $R_1$ and one colonizing bacterium $B_\alpha$ within any dilution cycle now reads as

$$\frac{dN_\alpha}{dt} = g_{\alpha 1} N_\alpha \, ; \quad \frac{dR_1}{dt} = -\frac{g_{\alpha 1}\, N_\alpha}{Y_{\alpha 1}}, \tag{S.50}$$

where $N_\alpha$ is the concentration of the microbe $B_\alpha$ and $Y_{\alpha 1}$ is the yield of the microbe $B_\alpha$ with respect to the resource $R_1$. Eq. (S.50) translates into the conservation law valid through a dilution cycle

$$N_\alpha + Y_{\alpha 1}\, R_1 = \text{const.} \tag{S.51}$$

In a steady state, the concentration of the resident should increase $D-$fold through a dilution cycle ($D$ being the dilution coefficient), so that

$$DN_\alpha = N_\alpha + Y_{\alpha 1} R_1, \tag{S.52}$$

where, in contrast with Eqs. (S.50) and (S.51), $N_\alpha$ denotes the steady state concentration of the microbe resident, meaning the value of the concentration with which $B_\alpha$ opens each dilution cycle.

When there are two sources $R_1, R_2$ present in the medium, and the top choice of $B_\alpha$ is, say, $R_1$, we get $N_\alpha \exp(g_{\alpha 1} T_1) = N_\alpha + Y_{\alpha 1} R_1$, $T_1$ being the depletion time of $R_1$ in the steady state, and

$$DN_\alpha = N_\alpha + Y_{\alpha 1} R_1 + Y_{\alpha 2} R_2. \tag{S.53}$$

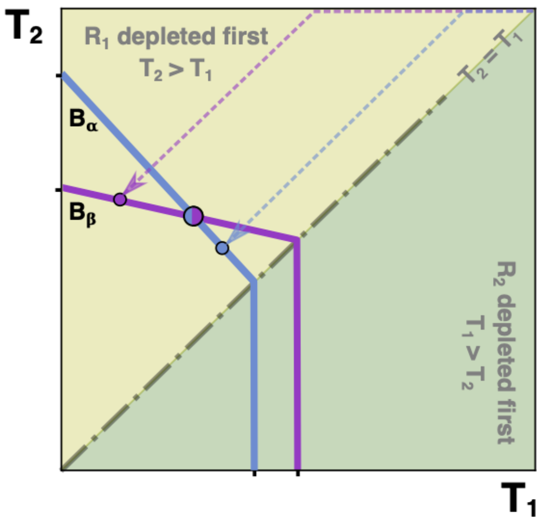

Figure S18: **Bistability in the environment of two sources open to invasion by two competing normal microbes.**

Together with the steady state condition in terms of the depletion times of the resources $T_1$ and $T_2$ which goes as $g_{\alpha 1}\, T_1 + g_{\alpha 2}\,(T_2 - T_1) = \log D$, that defines the depletion times as

$$T_1 = \frac{1}{g_{\alpha 1}} \log \frac{D\, Y_{\alpha 1}\, R_1 + Y_{\alpha 2}\, R_2}{Y_{\alpha 1}\, R_1 + Y_{\alpha 2}\, R_2}, \quad T_2 = T_1 + \frac{1}{g_{\alpha 2}} \log \frac{D\,(Y_{\alpha 1}\, R_1 + Y_{\alpha 2}\, R_2)}{D\, Y_{\alpha 1}\, R_1 + Y_{\alpha 2}\, R_2}. \tag{S.54}$$

This is where we have to reconsider the notion that, assuming the balanced nutrient supply, the time spent growing on the top choice is always the largest. Say, if the growth rates are close to each other, $D = 10$, $Y_{\alpha 1} = 0.1$, $Y_{\alpha 2} = 1$, $R_1 = R_2$, then $T_1/T_2 = 0.35\, g_{\alpha 2}/g_{\alpha 1}$, so that $T_1$ can be considerably less than $T_2$.

This makes a difference in two ways. First, the anomalous bacteria get a fair chance if they have a small yield on their top choice (unless the dilution coefficient is very large). At this point one can venture a prediction: the anomalous microbes that survive the competition with the normal ones have smaller yields on their top choice. Or, generalizing a statement made in the main text: **the evolutionary pressure encourages the microbes with the diauxie growth to arrange their preferential lists in such a way that either the growth rate on a source gets smaller or the yield gets larger down the list, the lower the rank of the source.** Second, even in the community of the normal microbes multistabilities can occur due to the differences in the yields. It is easily shown that the situation depicted in

Fig. S18 is realizable, when $D$ is not too large, if we allow yields to be different. Whenever the one-microbe line of $B_\alpha$ lands on its ZNGI at the point where $B_\beta$ cannot invade, and vice versa, the bistability can occur. Indeed, if $B_\beta$ is the first to invade, in the steady it consumes the resource $R_1$ favored by both microbes fast enough whenever it has small enough yield wrt $R_1$, so that $B_\alpha$ cannot invade. If $B_\alpha$ is the first to invade, it grows faster and makes the overall time shorter, so that $B_\beta$ cannot grow in the environment shaped by its competitor. Hence the steady state is chosen by chance: whoever is the first to invade, wins. All possible complexities can emerge when $D \sim 1/Y$.

We do not withhold the main conclusion concerning the optimization of complementarity even when there is a difference in the yield parameters. The microbe that has the best growth rate on the resource that is the last to disappear always has the advantage and thus can invade. This makes sorting by the top choice inevitable. However, the anomalous microbes might show up and, on rare occasions, become responsible for the low diversity states uninvadable by their normal counterparts.

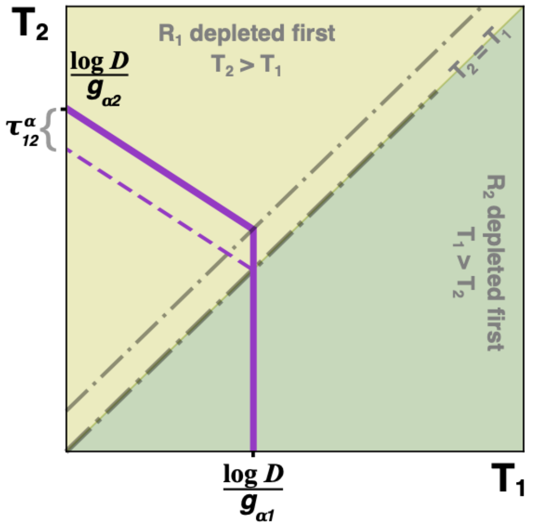

Figure S19: **Tilman-inspired diagram for one microbe, the effect of the lag.** The dashed lilac line represents the ZNGI of the microbe $B_\alpha$ with the top choice $R_1$ in absence of any lag. The solid line of the same color corresponds to the ZNGI with the lag modeled as a time period $\tau_{12}^{\alpha}$ of zero growth between the moment of complete consumption of the resource $R_1$ and that of starting the exponential growth on the resource $R_2$. There is no distinction between solid and dashed lines when the microbe grows on its top choice only (below the diagonal).

### H Taking lags into account

The simplest way to incorporate the lag phase into the model would be to ascribe a special value of the "lag time" to each diauxic shift event. Assume that, sensing that the nutrient currently in use is getting scarce, a microbe pauses in its growth and prepares the metabolic tools for another resource. The pause during which the species $B_\alpha$ switches from $R_1$ to $R_2$ we denote as $\tau_{12}^{(\alpha)}$.

The equation defining the microbe's ZNGI would take the following form:

$$g_{\alpha 1}T_1 + g_{\alpha 2}(T_2 - T_1 - \tau_{12}^{(\alpha)}) = \log\, D, \qquad \text{(S.55)}$$

$T_1, T_2$ being the depletion times of the resources $R_1$, $R_2$; $g_{\alpha 1}$ and $g_{\alpha 2}$ — the growth rates of the species $S_\alpha$ on each on those resources in their turn, $D$ — the dilution coefficient, as usual.

Fig. S20 shows how a lag of the simplest kind is treated by a Tilman-inspired diagram. In the Time plane, the diagonal $T_1 = T_2$ no longer separates the regions corresponding to the two different depletion orders. The boundary shifts up, exactly by the value of the lag time $\tau_{12}^{\alpha}$. The lifeline in the upper region where $T_2 > T_1$ and $R_1$ is the first to be depleted also shifts up, parallel to the lag-free version, and by the same amount $\tau_{12}^{\alpha}$.

In the main text we have made it clear that, in a mature community, the complementarity tends to become stronger, almost to the point of niche separation, so that (1) depletion times of different resources become close to each other; (2) the feeding strategy of the surviving community in a stable state is, most often, "to each its own top-choice", meaning that the main bulk of the biomass acquired by a species during a dilution step comes from its top-choice nutrient. In fact, the only thing in which the mature community of the species which are "specialists" (meaning that they specialize in using one and only nutrition source) differs from that of the "diauxic shifters", or "switchers" in this particular experimental setting is that, for the latter, the depletion times of the nutrients evolve to become almost equal. This was the conclusion based on the model that does not take lags into account. However, it is even more pronounced when we do take lags into account. It turns out, in a way, lags promote niche

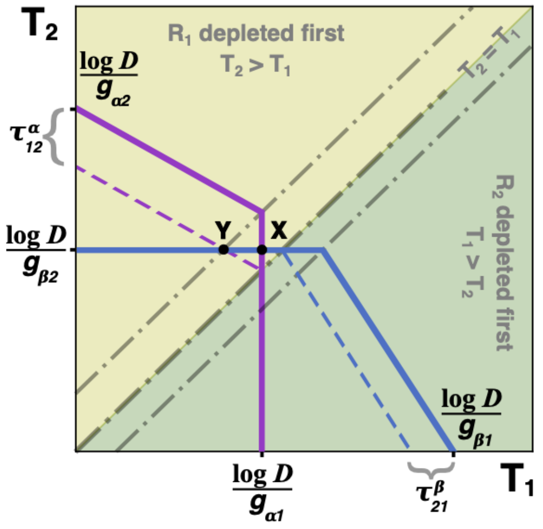

Figure S20: **Lags favour specializing.** The dashed segments of the ZNGI lines (lilac, for the microbe $B_\alpha$ and light blue, for $B_\beta$) represent the situation in absence of lags. If there had been no lags, both microbes would have been able to coexist at the point **Y**, with $B_\alpha$ having used both resources through a dilution cycle while $B_\beta$ growing on its top choice $R_2$. In presence of lags, the coexistence point switches to **X**. At that point, each microbe gets to grow on its own top choice resource only. The lag times are $\tau_{12}^{\alpha}$ for $B_\alpha$ (required to switch from the resource $R_1$ to the resource $R_2$) and $\tau_{21}^{\beta}$ for $B_\beta$ (required for switching from the resource $R_2$ to the resource $R_1$. Solid lines represent the ZNGI with the lags taken into account.

separation.

In Fig. S20 the point $X$ indicates the position of coexistence for two species $B_\alpha$ and $B_\beta$, lags taken into account. As it is obvious from the way the species' ZNGI lines go, $B_\alpha$ first consumes $R_1$ and then, after a lag time $\tau_{12}^{(\alpha)}$, switches to the resource $R_2$, while in the case of the species $B_\beta$, the order of preferences is reversed. The point $X$ is where the species coexist, driven to the "neutral band" by their lag times. At this steady state, through each dilution step, each of the microbe grows on its top-choice nutrient only. The lag condition drives the community (consisting, in this particular case, of two species) to a complete (virtual) niche separation. In absence of lags, the two species would have been able to coexist at the point $Y$, at which $B_\alpha$ consumes $R_1$, then $R_2$, while $B_\beta$ never gets a taste of $R_1$. If such were the case, the niche separation would have stopped halfway, as the lags had not been there to drive it on.

Regarding the question of a trade-off between the growth rate on a preferred nutrient versus the shortness of the lag phase[9, 10], it does play a role in the community forming.

In absence of such a trade-off, as is seen in Fig. S21, the ZNGI lines of the two species with the same preference order experience a parallel shift and otherwise coexist under the same external conditions (meaning the ratio $R_1/R_2$). On the other hand, if such a trade-off is present, it may even promote coexistence.

In Fig. S22 the species $B_\alpha$ chooses a large growth rate on its preferred nutrient $R_1$ at the expense of a huge lag time $\tau_{12}^{(\alpha)}$ while $B_\beta$ preferring the same nutrient opts for a smaller growth rate while keeping its lag reasonably small as well. With no lag and the same sets of growth rates, the species would not be able to coexist as $B_\alpha$ would win either way. However, as there is that trade-off, they coexist at the point $X$. (Note that in reality in the absence of that trade-off their growth rates might have been different from those defining the ZNGI slopes in the Fig. S22, so the notion that the trade-off promotes coexistence may be partly misleading.)

The presence of lags makes life harder for the anomalous species whose nutrition strategy is to grow with the higher rate on its second choice rather than on its first. While that may bring success under certain conditions (namely, considerable imbalance in nutrient supply, see

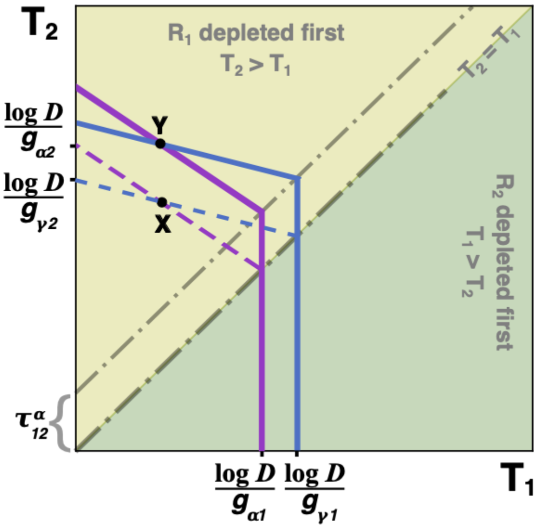

Figure S21: **In the absence of a trade-off (lag time vs growth rate). X** is the coexistence point of the species $B_\alpha$ and $B_\gamma$ sharing the same top choice $R_1$ in the absence of lags. If there is no trade-off between the growth rate and the length of the lag period, accounting for the lag time moves the point **X** up to the point **Y** by parallel translation. The conditions for the coexistence remain the same. (Dashed lines are for ZNGI in absence of the lags, solid lines account for the presence of the lags.)

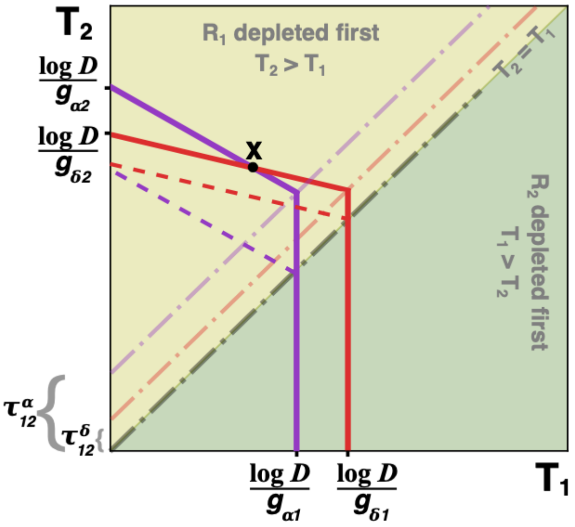

Figure S22: **When there is a trade-off (lag time vs growth rate)** In the presence of the trade-off, the bacterium $B_\alpha$ (lilac ZNGI) with the larger growth rate on its top choice $R_1$ is expected to exhibit longer period of the lag time when switching from $R_1$ to $R_2$. At the smaller values of the ratio $R_1/R_2$ the microbe $B_\delta$ (red ZNGI) that opts for the lesser growth rate on its top choice $R_1$ and shorter lag time gets an advantage. The coexistence is possible at the point **X**. If the microbes had the same growth rates with no lags (or with the lags, but in absence of the trade-off), $B_\alpha$ would have won the competition at any values of the ratio $R_1/R_2$.

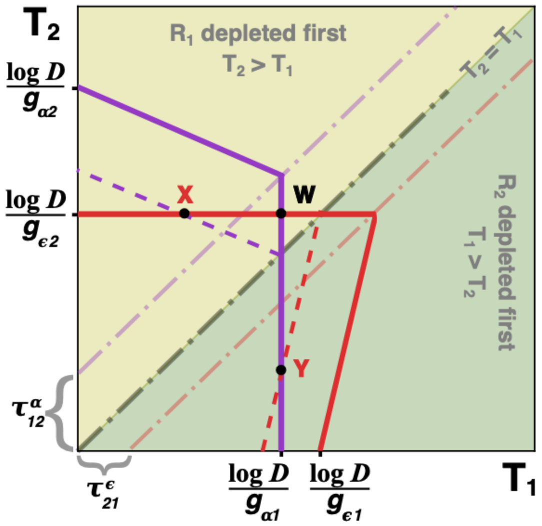

Figure S23: **Regular lag times lessen chances of survival for anomalous species.** In absence of lags, the anomalous species $B_\epsilon$ (the red ZNGI, dashed "lag-less" version) would have been able to coexist with the normal bacterium $B_\alpha$ (the lilac ZNGI, dashed "lag-less" version) at the point **X** in a stable way and at the point **Y** in an unstable way, allowing the anomalous species become a single uninvadable resident within a particular range of the values of $R_1/R_2$. The latter option becomes impossible in presence of lags (solid ZNGI lines). The only point of coexistence is now **W** at which the anomalous species does not get to use the resource $R_1$ at which it grows fastest.

the corresponding supplementary section), the advantage is easily lost when (regular) lag times come into light.

In Fig. S23 the red ZNGI corresponds to an anomalous species $B_\epsilon$ (that can be judged by the slope of its lifeline at $T_1 > T_2$). In a way, the species is a "superbug": it is not at all bad at competing for its top choice which is $R_2$, and on its second choice $R_1$ it grows even faster. The regular lag, however, rends the coexistence point $Y$ ineffective, so that under its most favourable depletion order the species cannot grow. It can still survive at the point $W$ if it is good enough on its top choice, but that would mean it should excel on both accounts while a normal species may be doing fine if it competes reasonably well just for its top choice. This would make the steady states containing anomalous species extremely rare after a considerable number of invasion events. The nature of possible trade-offs however could be finer on that particular point. There are indications[9, 10] that the trade-off goes between the growth rate on the top choice and the length of the lag phase preceding switching from the top choice to the second one. A wild guess would be that in some cases the threshold of "zero lag time" is trespassable, and a reason for some species being anomalous could be that they have, in fact, gone very far in their trade-offs and attained a negative lag time in switching from the nutrient they grow on best to the one on which they grow slower. If this were the case, there would have been no lag in the usual sense between the top choice resource and the second choice for the anomalous species. Can there exist a biological mechanism wild enough to support that wild guess?

In any case, the conclusion of this section is that the features of the mature steady state microbial community (as a result of the modeling done without taking the lag phenomenon into consideration) get even more pronounced due to the presence of lags.

##### H.1 Multistability induced by lags

Yet another type of multistability that can arise in the serial dilution setting is due to the presence of lags. The lag time is not a solely diauxic feature. If a bacterial culture is starved and is then introduced to a resource, there is usually some time before it can resume growing. Such a regrowth lag is not of a diauxic nature and can be experienced by a co-utilizer [11]. For all we know, its length may be affected by the previous circumstances, i. e. whether the last resource the microbe has grown upon is the same as the one to which it is introduced after a period of starving, or different. In this section we try and take the regrowth lags into account as well as the diauxic lags.

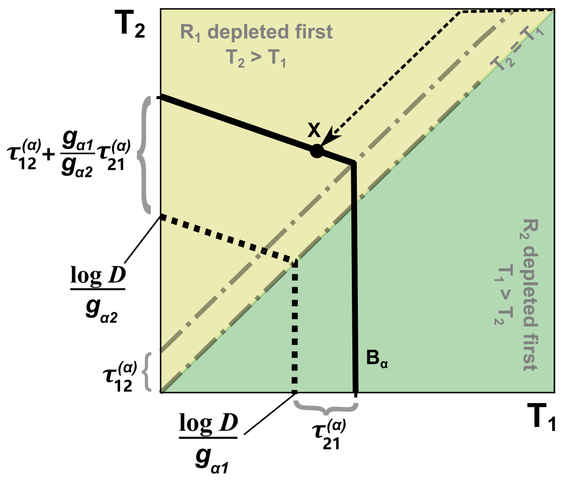

Figure S24: **ZNGI with the effects of both regrowth and diauxie lags indicated.** The dashed segments of the ZNGIs stand for the situation of $B_\alpha$'s monoculture in absence of lags. With both lags ($\tau_{12}^{(\alpha)}$ represents the diauxie lag, and $\tau_{21}^{(\alpha)}$ the regrowth lag), the ZNGIs become what is shown by the solid black lines. The steady state is represented by point X.

Fig.(S24) shows a zero net growth isocline (ZNGI) of a microbe $B_\alpha$ in the Time plane $(T_1, T_2)$: the axes, as usual, being the depletion times of the resources $R_1$ and $R_2$ present in the environment. The top choice of the microbe is the resource $R_1$. When a dilution cycle starts, the (starved) monoculture of the microbe $B_\alpha$ is introduced to the environment containing both $R_1$ and $R_2$. The bacterium experiences a regrowth lag $\tau_{21}^{(\alpha)}$ and then starts consuming $R_1$. Upon depletion of the resource and the diauxic lag period of the length $\tau_{12}^{(\alpha)}$ the microbe switches to the resource $R_2$, consumes it and is starved for the rest of the dilution cycle. (At the very end, it gets diluted $D-$fold and introduced to the similar environment as at the beginning of the previous cycle.) Note that we use the designation $\tau_{21}^{(\alpha)}$ for the regrowth lag time to indicate that the length of the lag may depend on the history of consumption. The slope of the ZNGI line (where it is inclined wrt both axes) in this model does not depend on the lag values, but the height of the point where it hits the $T_2$ axis or the degree to which it is raised above the diagonal does. So, in theory, the lag time as a controlling parameter could bring the ZNGI lines of different microbes up or down, make them intersect when they otherwise wouldn't and so on. This, however, could result in a multistability (bistability, if only two sources are available for consumption).

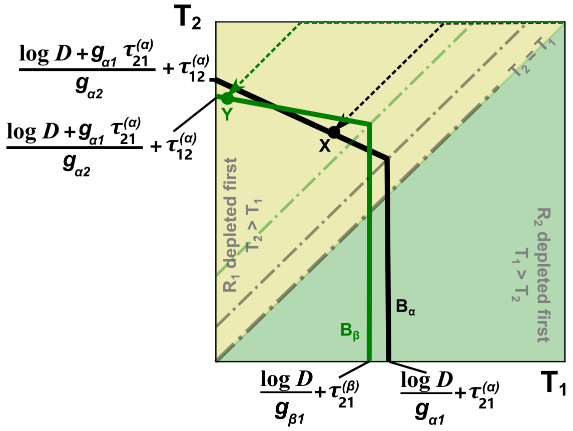

Figure S25: **Lags prompting bistability.** With the presence of both regrowth lag and diauxie lag, it is possible that species $B_\alpha$ (ZNGI shown by black solid lines) and $B_\beta$ (green solid lines) with the same preference order can mutually exclude each other. The monoculture steady states of these two species correspond to point X and Y.

In Fig.(S25) the microbe $B_\alpha$ with the steeper slope of the (black) ZNGI has its monoculture steady state situated at the point $(T_1^{(\alpha)}, T_2^{(\alpha)})$ where the microbe $B_\beta$ (green ZNGI) cannot grow, and vice versa: if $B_\beta$ is the first to establish its monoculture steady state, $B_\alpha$ would not be able to invade. This can be made possible in two ways. First, if $B_\alpha$ grows faster than $B_\beta$ on their top choice, i. e. $g_{\alpha 1} > g_{\beta 1}$, its one-microbe line would normally land on its ZNGI to the right on that of $B_\beta$ (see the subsection **??** of the Supplementary Text). If however the regrowth lag of $B_\alpha$ is longer than that of $B_\beta$, i. e. $\tau_{21}^{(\alpha)} > \tau_{21}^{(\beta)}$ (thus moving the former's one microbe line farther to the right), such a disposition is realizable. Second, even if $g_{\alpha 1} < g_{\beta 1}$, but $g_{\alpha 1}/g_{\alpha 2} > g_{\beta 1}/g_{\beta 2}$ and the diauxie lag of $B_\beta$ is larger than that of $B_\alpha$, so that $\tau_{12}^{(\alpha)} < \tau_{12}^{(\beta)}$, the intersection may be possible and, as, in this case, the one microbe line of $B_\alpha$ normally lands to the right of that of $B_\beta$ (if the regrowth lags don't interfere), such a disposition may, again, be realizable.

In Barthe et al. the authors hint that the lag times can actually be controlled in bacteria[1]. If this can be indeed managed, then we are presented with a thrilling possibility to construct complex microbe communities at will, choosing winners or multistable scenarios by manipulating solely the lag parameters.

Now we proceed to deriving a formal criterion that decides the fate of such bistability. We assume $g_{\alpha 1}/g_{\alpha 2} > g_{\beta 1}/g_{\beta 2}$, like it is shown in Fig.S25 (the above ratios, or, to be precise, the ratios $(g_{\alpha 1} - g_{\alpha 2})/g_{\alpha 2} > (g_{\beta 1} - g_{\beta 2})/g_{\beta 2}$ controlling the slopes of the upper parts of the ZNGI lines).

The steady state depletion times for the $B_\alpha$ monoculture are

$$\begin{cases} T_1^{(\alpha)} = \dfrac{1}{g_{\alpha 1}} \log\left(\dfrac{D\,R_1 + R_2}{R_1 + R_2}\right) + \tau_{21}^{(\alpha)} \\ T_2^{(\alpha)} = T_1^{(\alpha)} + \tau_{12}^{(\alpha)} + \dfrac{1}{g_{\alpha 2}} \log\left(\dfrac{D\,(R_1 + R_2)}{D\,R_1 + R_2}\right). \end{cases} \tag{S.56}$$

If $B_\beta$ cannot invade, that means its multiplication coefficient with those depletion times is less than $D$, i. e.

$$g_{\beta 1}\,(T_1^{(\alpha)} - \tau_{21}^{(\beta)}) + g_{\beta 2}\,(T_2^{(\alpha)} - T_1^{(\alpha)} - \tau_{12}^{(\beta)}) < \log D, \tag{S.57}$$

which, combined with (S.56), gives

$$\frac{g_{\alpha 1}\Big((g_{\beta 2} - g_{\alpha 2})\,\log D + g_{\alpha 2}\,(g_{\beta 1}\,\Delta\tau_{21} + g_{\beta 2}\,\Delta\tau_{12})\Big)}{g_{\alpha 1}\,g_{\beta 2} - g_{\alpha 2}\,g_{\beta 1}} < \log\left(\frac{DR_1 + R_2}{R_1 + R_2}\right), \tag{S.58}$$

where $\Delta\tau_{21} = \tau_{21}^{(\alpha)} - \tau_{21}^{(\beta)}$ and $\Delta\tau_{12} = \tau_{12}^{(\alpha)} - \tau_{12}^{(\beta)}$.

Note that, for (S.58) to be plausible, the left-hand side of the equation should be less than $\log D$. That implies

$$(g_{\beta 1} - g_{\alpha 1}) \log D + g_{\alpha 1} (g_{\beta 1} \Delta\tau_{21} + g_{\beta 2} \Delta\tau_{12}) < 0. \tag{S.59}$$

If, on the other hand, the monoculture of $B_\beta$ was the first to establish itself in the environment, the depletion times of the steady state would be

$$\begin{cases} T_1^{(\beta)} = \dfrac{1}{g_{\beta 1}} \log\Big(\dfrac{D\,R_1 + R_2}{R_1 + R_2}\Big) + \tau_{21}^{(\beta)} \\ T_2^{(\beta)} = T_1^{(\beta)} + \tau_{12}^{(\beta)} + \dfrac{1}{g_{\beta 2}} \log\Big(\dfrac{D\,(R_1 + R_2)}{D\,R_1 + R_2}\Big), \end{cases} \tag{S.60}$$

and $B_\alpha$ would not be able to invade if

$$g_{\alpha 1} (T_1^{(\beta)} - \tau_{21}^{(\alpha)}) + g_{\alpha 2} (T_2^{(\beta)} - T_1^{(\beta)} - \tau_{12}^{(\alpha)}) < \log D, \tag{S.61}$$

which, combined in its turn with (S.60), leads to

$$\frac{g_{\beta 1} \Big((g_{\beta 2} - g_{\alpha 2}) \log D + g_{\beta 2} (g_{\alpha 1} \Delta\tau_{21} + g_{\alpha 2} \Delta\tau_{12})\Big)}{g_{\alpha 1}\, g_{\beta 2} - g_{\alpha 2}\, g_{\beta 1}} > \log\Big(\frac{DR_1 + R_2}{R_1 + R_2}\Big), \tag{S.62}$$

The above can be plausible only if the left-hand side of Eq. (S.62) is positive. That spells as

$$(g_{\beta 2} - g_{\alpha 2}) \log D + g_{\beta 2} (g_{\alpha 1} \Delta\tau_{21} + g_{\alpha 2} \Delta\tau_{12}) > 0. \tag{S.63}$$

To be able to match (S.58) with (S.62), one must see to it that the left-hand side of the latter is greater than the left-hand side of the former, and that reduces to

$$-(g_{\alpha 1} - g_{\beta 1}) (g_{\alpha 2} - g_{\beta 2}) \log D + g_{\alpha 1}\, g_{\beta 1} (g_{\alpha 2} - g_{\beta 2}) \Delta\tau_{21} + g_{\alpha 2}\, g_{\beta 2} (g_{\alpha 1} - g_{\beta 1}) \Delta\tau_{12} < 0. \tag{S.64}$$

The Eqs (S.59), (S.63) and, most importantly, (S.64) form the set of conditions under which there exists a range of the $R_1/R_2$ ratio values within which bistability can occur. Before we find out the formal range of that ratio in case it exists, let us look closer at the Eq. S.64. First, in absence of lags (or if the corresponding lags are equal to each other, $\tau_{21}^{(\alpha)} = \tau_{21}^{(\beta)}$ and $\tau_{12}^{(\alpha)} = \tau_{12}^{(\beta)}$), the Eqs (S.59), (S.63) and (S.64) are obviously incompatible, so that there may be no bistability in this case. If however it can be assumed that $|\Delta\tau_{12}| \ll |\Delta\tau_{21}|$, (S.64) reduces to

$$(g_{\alpha 2} - g_{\beta 2})(-(g_{\alpha 1} - g_{\beta 1}) \log D + g_{\alpha 1}\, g_{\beta 1} \Delta\tau_{21}) < 0 \tag{S.65}$$

If $g_{\alpha 1} > g_{\beta 1}$ and $g_{\alpha 2} < g_{\beta 2}$, then, just as we have discussed in the beginning of this subsection, to get a chance to obtain bistability here we should have $\Delta\tau_{21} > 0$; moreover, as (S.59) in this case actually coincides with (S.65) and (S.63) provides no additional restrictions, we obtain

$$\Delta\tau_{21} > \frac{g_{\alpha 1} - g_{\beta 1}}{g_{\alpha 1}\, g_{\alpha 2}} \log D \tag{S.66}$$

If, on the contrary, the difference in the regrowth lags of both microbes is negligible compared to that of their diauxic lag times, $|\Delta\tau_{12}| \gg |\Delta\tau_{21}|$, then (S.64) becomes

$$(g_{\alpha 1} - g_{\beta 1})(-(g_{\alpha 2} - g_{\beta 2}) \log D + g_{\alpha 2}\, g_{\beta 2} \Delta\tau_{12}) < 0 \tag{S.67}$$

If $B_\alpha$ loses to $B_\beta$ on both accounts in terms of growth rates, $\Delta\tau_{12}$ cannot be positive due to the condition (S.59), and we must have

$$-\frac{g_{\beta 2} - g_{\alpha 2}}{g_{\alpha 2}\, g_{\beta 2}} \log D < \Delta\tau_{12} < -\frac{g_{\beta 1} - g_{\alpha 1}}{g_{\alpha 1}\, g_{\beta 2}} \log D, \tag{S.68}$$

which works out well under our condition $g_{\alpha1}/g_{\alpha2} > g_{\beta1}/g_{\beta2}$ assumed in the begining for the sake of definitiveness.

Note that, in the case $g_{\alpha1} > g_{\beta1}$ Eqs (S.63) and (S.68) become incompatible, so that in this case the bistability can be provided only by manipulating the difference of the regrowth lag times $\Delta\tau_{21}$.

Assume that the conditions (S.59), (S.63) and (S.64) are all satisfied and proceed to determine the range of $R_1/R_2$ ratio that favours bistability. Introduce the notations

$$\begin{aligned} x &= \frac{g_{\alpha1}\,(g_{\beta2} - g_{\alpha2})}{g_{\alpha1}\,g_{\beta2} - g_{\alpha2}\,g_{\beta1}} \\ y &= \frac{g_{\alpha1}\,g_{\alpha2}\,(g_{\beta1}\,\Delta\tau_{21} + g_{\beta2}\,\Delta\tau_{12})}{g_{\alpha1}\,g_{\beta2} - g_{\alpha2}\,g_{\beta1}}. \end{aligned} \tag{S.69}$$

Eq. (S.58) then takes the form

$$D^x \exp(y) < \frac{DR_1 + R_2}{R_1 + R_2}. \tag{S.70}$$

As we have made sure that $D^x \exp(y) < D$, the above is equivalent to

$$\frac{R_1}{R_2} > \frac{D^x \exp(y) - 1}{D - D^x \exp(y)}. \tag{S.71}$$

Again, introduce the notations:

$$\begin{aligned} u &= \frac{g_{\beta1}\,(g_{\beta2} - g_{\alpha2})}{g_{\alpha1}\,g_{\beta2} - g_{\alpha2}\,g_{\beta1}} \\ v &= \frac{g_{\beta1}\,g_{\beta2}\,(g_{\alpha1}\,\Delta\tau_{21} + g_{\alpha2}\,\Delta\tau_{12})}{g_{\alpha1}\,g_{\beta2} - g_{\alpha2}\,g_{\beta1}}. \end{aligned} \tag{S.72}$$

The left-hand side of Eq. (S.62) will than read as $D^u \exp(v)$. There is a possibility that $D^u \exp(v) > D$, i. e.

$$(g_{\beta1} - g_{\alpha1})\log D + g_{\beta1}\,(g_{\alpha1}\,\Delta\tau_{21} + g_{\alpha2}\,\Delta\tau_{12}) > 0, \tag{S.73}$$

then there are no additional restrictions coming from the fact that $g_\alpha$ cannot invade $B_\beta-$monoculture, so that (S.71) defines the region of $R_1/R_2$ favouring the bistability phenomenon. If however (S.73) does not hold, and the opposite is true, then we get yet another condition

$$D^u \exp(v) > \frac{DR_1 + R_2}{R_1 + R_2}, \tag{S.74}$$

which, combined with (S.71), results in the final bistability condition

$$\frac{D^x \exp(y) - 1}{D - D^x \exp(y)} < \frac{R_1}{R_2} < \frac{D^u \exp(v) - 1}{D - D^u \exp(v)}, \tag{S.75}$$

with $x, y$ given by (S.69) and $u, v$ by (S.74).

To conclude, we note that if, say, $g_{\alpha1} > g_{\beta1}$ and $g_{\alpha2} < g_{\beta2}$, the lag times and the dilution coefficient can be arranged so that the competition between the two species would rend any of them as a sole winner, or result in their co-existence, or even display the tendency towards bistability.

### I Representing the co-utilizing microbes in the time plane

Earlier (in the main text) we have claimed that the geometrical approach we have developed is useful whichever metabolic strategy is chosen by the microbes in the environment subject to

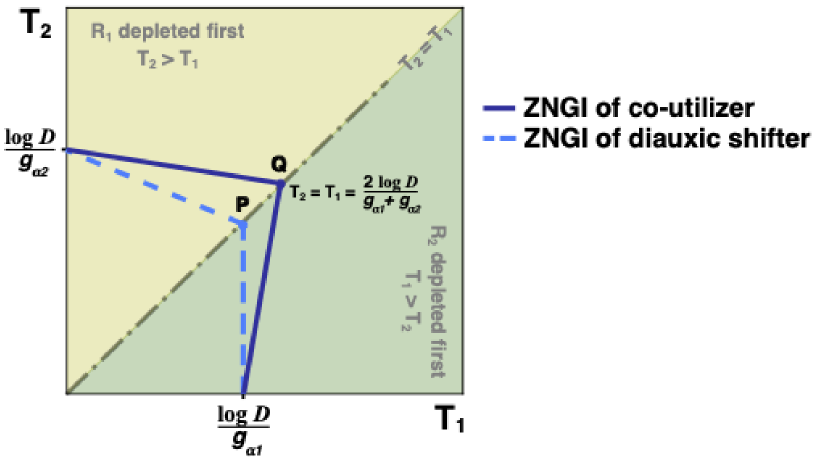

Figure S26: **Zero net growth isoclines: co-utiliser vs diauxic shifter in the Time plane.** The co-utilising microbe (violet ZNGI) and the diauxie shifting microbe (dashed blue ZNGI, top choice $R_1$) technically have the same growth rates on both resources $R_1$ and $R_2$. However, in absence of lags the diauxie shifter's ZNGI is located inside the Time plane segment bounded by the co-utiliser's zero net growth isocline, so that at any values of the ratio $R_1/R_2$ the former would win the competition. One of the key indicators of how a microbe fares at a roughly balanced nutrient supply is the position of the point at which the microbe's ZNGI intersects with the diagonal $T_1 = T_2$. It can be noted that the point $P$ associated with the diauxie-shifting microbe is closer to the origin than the point $Q$ associated with the co-utiliser.

serial dilution. Here we suggest the way to treat the co-utilizing microbes in the context of the serial dilution set-up. As in the case with the metabolic strategy involving diauxie, we assume that each dilution cycle starts with the initial abundance of resources sufficiently high to ensure exponential growth of the resident microbes. We also assume for simplicity that the co-utilising microbes given $n$ resources allocate to each one $1/n$ fraction if its metabolic engines, so that a microbe $B_\alpha$ grows on the resources $R_1, R_2, ..., R_n$ with the rate $(g_{\alpha 1} + g_{\alpha_2} + ... + g_{\alpha n})/n$. That would mean that, in the case of the environment containing two resources $R_1$, $R_2$, the microbe $B_\alpha$ that happens to deplete first $R_1$ and then $R_2$ (utilising both while they are available), for the time period $T_1$ will grow with the rate $(g_{\alpha 1} + g_{\alpha 2})/2$ and then for the time period $T_2 - T_1$ with the rate $g_{\alpha 2}$. Hence the zero net-growth isocline (ZNGI) for this particular depletion order will have the equation

$$\frac{g_{\alpha 1} + g_{\alpha 2}}{2}\, T_1 + g_{\alpha 2}\,(T_2 - T_1) = \log\, D, \tag{S.76}$$

where $D$ is, as usual, the dilution coefficient. Should the depletion order happen to be reversed, so that the depletion times become such that $T_1 > T_2$, $B_\alpha$ would then grow first for the time period $T_2$ with the rate $(g_{\alpha 1} + g_{\alpha 2})/2$ and then, for the time period $T_1 - T_2$, with the rate $g_{\alpha 1}$. That depletion order corresponds to the part of the ZNGI having the equation

$$\frac{g_{\alpha 1} + g_{\alpha 2}}{2}\, T_2 + g_{\alpha 1}\,(T_1 - T_2) = \log\, D. \tag{S.77}$$

When both resources are depleted at the same time, one gets $T_1 = T_2 = \log\, D/((g_{\alpha 1} + g_{\alpha 2})/2)$, which is different from the case of the diauxic shifters. For the latter, when the depletion times are equal, they are also equal to $\log\, D/\max(g_1, g_2)$, i. e. (for the normal species) with the maximum of the two growth rates on two different nutrients in the denominator rather than the average growth rate. This difference implies that, in the case of the co-utilizing microbes, the growth rate "on the top choice" (which translates simply into the largest growth rate, as the co-utilizing microbes use all the resources simultaneously) may cease to be the main selection parameter.

In Fig. S26 we have shown the ZNGI line for the co-utiliser $B_\alpha$ (solid line) and (dashed line) the way the same isocline would go if $B_\alpha$ were a diauxic shifter, with the same values of growth rates, and the preference order first $R_1$, then $R_2$. The figure may be misleading in two ways: (1) the co-utilisers metabolic strategy is modeled too crudely; (2) in the case of the diauxic shifter, it does not show any lag. Still, the message is clear: as long as the lag is smaller than some critical lag value determined by the growth rates and the dilution rate, in the case of the balanced (and regular) nutrient supply it is more reasonable to opt for a diauxic shift than for co-utilising.

We do not go into further details concerning the (fascinating) differences between the two metabolic strategies (easily brought to light by the graphic approach), as those are clearly beyond the scope of this particular work.

#### J Coexistence of the diauxic species having the same preferential orders

While the species with the same preferential order with regard to the carbon sources are naturally pressed harder by the competitive exclusion principle, they can still co-exist when the external conditions are right. The advantage of our model being its simplicity, the set of the external conditions in our case boils down to that of the relative abundances of the initial resource concentrations, e. g. $R_1/R_2$ if there are two resources available, and the value of the dilution coefficient $D$. In this subsection we will obtain the conditions allowing for the coexistence of two species $B_\alpha$ and $B_\beta$, both having the resource $R_1$ as their top choice, and, upon its depletion, switching to the resource $R_2$ in a diauxie manner. We will also try to include the diauxic lags into the consideration as well as point out the conditions under which their presence may be neglected. Using the data provided by Barthe et al. [1], we will make a testable prediction specifying, for various values of the dilution coefficient $D$, the region of $R_1/R_2$ favourable for the coexistence of the strains $E22$ and $M1/5$ of the bacterium *E. coli* growing on a mixture of glucose and xylose in the serial dilution setting.

Consider microbes $B_\alpha$ and $B_\beta$ able to grow on the carbon sources $R_1$ and $R_2$ in a diauxic manner with the growth rates $g_{\alpha 1} > g_{\beta 1}$ on the first resource and $g_{\alpha 2} < g_{\beta 2}$ on the second. When $R_1$ and $R_2$ are both available, the microbes start with $R_1$ and, when it is consumed, after the lag periods $\tau_{12}^{(\alpha)}$ and $\tau_{12}^{(\beta)}$ respectively start consuming $R_2$. Presumably at the beginning of the next dilution cycle both microbes demonstrate a regrowth lag when being intoduced again to $R_1$, $\tau_{21}^{(\alpha)}$ and $\tau_{21}^{(\beta)}$ respectively. If the microbes share the environment, coexisting in a steady state, then their intial concentrations get multiplied by the factor $D$ till the end of each dilution cycle. Due to the exponential nature of their growth, we get the equations

$$\begin{cases} g_{\alpha 1}\,(T_1 - \tau_{21}^{(\alpha)}) + g_{\alpha 2}\,(T_2 - T_1 - \tau_{12}^{(\alpha)}) = \log D \\ g_{\beta 1}\,(T_1 - \tau_{21}^{(\beta)}) + g_{\alpha 2}\,(T_2 - T_1 - \tau_{12}^{(\beta)}) = \log D, \end{cases} \tag{S.78}$$

where $T_1$ and $T_2$ are the depletion times of the resources $R_1$ and $R_2$ respectively, as usual. We get

$$T_1 = \frac{(g_{\beta 2} - g_{\alpha 2})\,\log D + g_{\beta 2}\,g_{\alpha 1}\,\tau_{21}^{(\alpha)} - g_{\alpha 2}\,g_{\beta 1}\,\tau_{21}^{(\beta)} + g_{\alpha 2}\,g_{\beta 2}(\tau_{12}^{(\alpha)} - \tau_{12}^{(\beta)})}{g_{\alpha 1}\,g_{\beta 2} - g_{\alpha 2}\,g_{\beta 1}} \tag{S.79}$$

and

$$\begin{aligned} T_2 = \frac{1}{g_{\alpha 1}\,g_{\beta 2} - g_{\alpha 2}\,g_{\beta 1}}\,\Big(&(g_{\alpha 1} - g_{\alpha 2} - g_{\beta 1} + g_{\beta 2})\,\log D + \\ &+ (g_{\alpha 1} - g_{\alpha 2})\,(g_{\beta 1}\,\tau_{21}^{(\beta)} + g_{\beta 2}\,\tau_{12}^{(\beta)}) - (g_{\beta 1} - g_{\beta 2})\,(g_{\alpha 1}\,\tau_{21}^{(\alpha)} + g_{\alpha 2}\,\tau_{12}^{(\alpha)})\Big) \end{aligned} \tag{S.80}$$

In order to obtain the coexistence conditions for $B_\alpha$ and $B_\beta$ we determine in a formal way the steady state initial concentrations $N_\alpha$, $N_\beta$ of the microbes assuming they share the environment and then find at which values of the ratio $R_1/R_2$ both $N_\alpha$ and $N_\beta$ thus obtained are positive. (The alternative approach is to use the Tilman-inspired diagrams, and we will discuss it later in that same subsection as well.)

Denote $\Delta T_1^{(\alpha)} = T_1 - \tau_{21}^{(\alpha)}$ and $\Delta T_1^{(\beta)} = T_1 - \tau_{21}^{(\beta)}$ — the time periods during which $B_\alpha$ and $B_\beta$ respectfully grow on $R_1$ until it is finally depleted. Then, at the moment when the first resource is consumed, the biomass conservation law would have it

$$N_\alpha \exp\left(g_{\alpha 1}\,\Delta T_1^{(\alpha)}\right) + N_\beta \exp\left(g_{\beta 1}\,\Delta T_1^{(\beta)}\right) = R_1 + N_\alpha + N_\beta. \tag{S.81}$$

The steady state condition (or the same conservation law at the point when both resources are consumed), on the other hand, requires

$$D\,N_\alpha + D\,N_\beta = R_1 + R_2 + N_\alpha + N_\beta. \tag{S.82}$$

Combining (S.81) and (S.82) we get

$$\begin{cases} N_\alpha = \dfrac{\Big(D - \exp\left(g_{\beta 1}\,\Delta T_1^{(\beta)}\right)\Big)\,R_1 - \Big(\exp\left(g_{\beta 1}\,\Delta T_1^{(\beta)}\right) - 1\Big)\,R_2}{(D-1)\Big(\exp\left(g_{\alpha 1}\,\Delta T_1^{(\alpha)}\right) - \exp\left(g_{\beta 1}\,\Delta T_1^{(\beta)}\right)\Big)} \\[2ex] N_\beta = \dfrac{\Big(\exp\left(g_{\alpha 1}\,\Delta T_1^{(\alpha)}\right) - 1\Big)\,R_2 - \Big(D - \exp\left(g_{\alpha 1}\,\Delta T_1^{(\alpha)}\right)\Big)\,R_1}{(D-1)\Big(\exp\left(g_{\alpha 1}\,\Delta T_1^{(\alpha)}\right) - \exp\left(g_{\beta 1}\,\Delta T_1^{(\beta)}\right)\Big)} \end{cases} \tag{S.83}$$

It is required that $N_\alpha > 0$, $N_\beta > 0$, so that the region of the values of the ratio $R_1/R_2$ favorable to the coexistence of both species is restricted to

$$\frac{\exp\left(g_{\beta 1}\,\Delta T_1^{(\beta)}\right) - 1}{D - \exp\left(g_{\beta 1}\,\Delta T_1^{(\beta)}\right)} < \frac{R_1}{R_2} < \frac{\exp\left(g_{\alpha 1}\,\Delta T_1^{(\alpha)}\right) - 1}{D - \exp\left(g_{\alpha 1}\,\Delta T_1^{(\alpha)}\right)} \tag{S.84}$$

The explicit expressions for $\Delta T_1^{(\alpha)} = T_1 - \tau_{21}^{(\alpha)}$ and $\Delta T_1^{(\beta)} = T_1 - \tau_{21}^{(\beta)}$ are

$$\Delta T_1^{(\alpha)} = \frac{(g_{\beta 2} - g_{\alpha 2})\,\log D + g_{\beta 1}\,g_{\alpha 2}\,(\tau_{21}^{(\alpha)} - \tau_{21}^{(\beta)}) + g_{\alpha 2}\,g_{\beta 2}\,(\tau_{12}^{(\alpha)} - \tau(\beta)_{12})}{g_{\alpha 1}\,g_{\beta 2} - g_{\alpha 2}\,g_{\beta 1}} \tag{S.85}$$

and

$$\Delta T_1^{(\beta)} = \frac{(g_{\beta 2} - g_{\alpha 2})\,\log D + g_{\beta 2}\,g_{\alpha 1}\,(\tau_{21}^{(\alpha)} - \tau_{21}^{(\beta)}) + g_{\alpha 2}\,g_{\beta 2}\,(\tau_{12}^{(\alpha)} - \tau(\beta)_{12})}{g_{\alpha 1}\,g_{\beta 2} - g_{\alpha 2}\,g_{\beta 1}} \tag{S.86}$$

From (S.85) and (S.86) it is obvious that when the difference in the lag times for the two microbes is small enough (or, in the other words, the dilution factor $D$ is large enough), so that

$$\log D \gg \frac{g_{\beta 2}\,g_{\alpha 1}\,\Delta\tau_{21} + g_{\alpha 2}\,g_{\beta 2}\Delta\tau_{12}}{g_{\beta 2} - g_{\alpha 2}} \tag{S.87}$$

($\Delta\tau_{21}$ and $\Delta\tau_{12}$ here being the differences in the corresponding lags for two microbes), then the lags in (S.84) can be neglected.

For two strains of Escherichia coli, $B_\alpha = M1/5$ and $B_\beta = E22$, the growth rates on $R_1 =$ glucose and $R_2 =$ xylose given by [1] are $g_{\alpha 1} = 0.9h^{-1}$, $g_{\alpha 2} = 0.35h^{-1}$ and $g_{\beta 1} = 0.75h^{-1}$, $g_{\beta 2} = 0.65h^{-1}$ respectively. Even if the relevant values of $\Delta\tau$ were of the order of 0.5 hour, we would have the right hand side of the Eq. (S.87) $\sim 0.4 - -0.5$, which is about one tenth of $\log D$ at $D = 100$. Judging by the data exposed by [1] however the lag times themselves are less or equal to 0.5 hour, so that the difference between them must be even less. This is where we can hope to lose nothing much by neglecting the terms associated with the lag times $\tau$ in the nominator of (S.84). Bearing this in mind, we transform (S.84) into

$$\frac{D^x - 1}{D - D^x} < \frac{R_1}{R_2} < \frac{D^y - 1}{D - D^y} \tag{S.88}$$

with

$$x = \frac{g_{\beta 1}\left(g_{\beta 2} - g_{\alpha 2}\right)}{g_{\alpha 1} g_{\beta 2} - g_{\beta 1} g_{\alpha 2}} \text{ and } y = \frac{g_{\alpha 1}\left(g_{\beta 2} - g_{\alpha 2}\right)}{g_{\alpha 1} g_{\beta 2} - g_{\beta 1} g_{\alpha 2}}.$$

The condition (S.88) is relevant whenever the corresponding lag times of the co-existing microbes are close enough to each other, as they tend to mutually cancel each other. In our particular case, with $B_\alpha$ and $B_\beta$ being the $M1/5$ and $E22$ strains of E-coli, this boils down to

$$\frac{D^{0.70} - 1}{D - D^{0.70}} < \frac{R_1}{R_2} < \frac{D^{0.84} - 1}{D - D^{0.84}} \tag{S.89}$$

To make our message more practical, we calculate the range of the ratio of the glucose and xylose concentrations favorable for the coexistence of the two *E. coli* strains ($B_\alpha = M1/5$ an $B_\beta = E22$) for the values of the dilution factor $D = 10, 100, 1000$ *and* $10000$. The ranges are:

- for $\mathbf{D} = \mathbf{10}$ it's $0.80 < \dfrac{R_1}{R_2} < 1.92$;
- for $\mathbf{D} = \mathbf{100}$ it's $0.32 < \dfrac{R_1}{R_2} < 0.90$;
- for $\mathbf{D} = \mathbf{1000}$ it's $0.14 < \dfrac{R_1}{R_2} < 0.49$;
- for $\mathbf{D} = \mathbf{10000}$ it's $0.07 < \dfrac{R_1}{R_2} < 0.19$.

With the increase of the dilution factor $D$, the interval of coexistence gets shorter as the number of generations needed to obtain the fitting multiplication factor becomes larger and the differences tend to accumulate with generations.

The other way to derive the conditions (S.84) and (S.88) is to consider the diagram showing the Zero Net Growth isoclines of both microbes in the time plane (the coordinate axes being the depletion times of the carbon sources), in the spirit of Tilman's mechanistic approach. The one-microbe lines of the species (i. e. the dynamic trajectories of the monoculture states) land on the isoclines at the points corresponding to the steady states of the monocultures. The depletion times $T_1^{(\alpha)}$ and $T_1^{(\beta)}$ pertaining to the steady state of the monocultures $B_\alpha$ and $B_\beta$ can be obtained in the same way as described above, from the biomass conservation laws: namely, when the first resource is depleted, one writes $N_\alpha \exp\left(g_{\alpha 1}(T_1^{(\alpha)} - \tau_\alpha^{21})\right) = R_1 + N_\alpha$ (for the monoculture of the microbe $B_\alpha$) and $N_\beta \exp\left(g_{\beta 1}(T_1^{(\beta)} - \tau_\beta^{21})\right) = R_1 + N_\beta$ (for the monoculture of the microbe $B_\beta$). Coupled with the biomass conservation law applied at the point when both resources are consumed and the steady state condition requiring that a microbe should multiply its population by the factor $D$ at the end of the dilution cycle, $D\,N_\alpha = R_1 + R_2 + N_\alpha$ (or $D\,N_\beta = R_1 + R_2 + N_\beta$, respectfully), one obtains the monoculture depletion times as

$$T_1^{(\alpha)} = \frac{1}{g_{\alpha 1}} \log\left(\frac{D R_1 + R_2}{R_1 + R_2}\right) + \tau_\alpha^{21} \tag{S.90}$$

and

$$T_1^{(\beta)} = \frac{1}{g_{\beta 1}} \log\left(\frac{D R_1 + R_2}{R_1 + R_2}\right) + \tau_\beta^{21} \tag{S.91}$$

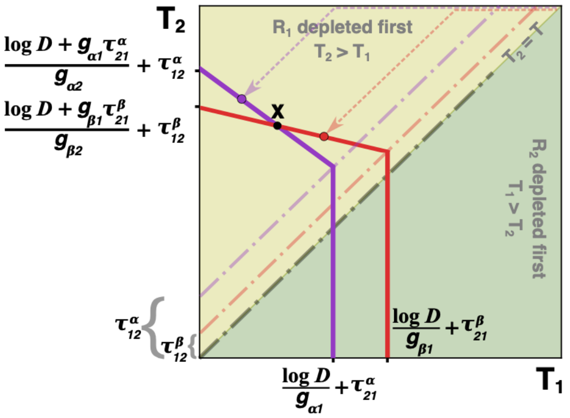

Figure S27: **Coexistence of the diauxic species having the same preferential orders.** The two *E. coli* species, with their non-zero initial and regrowth lags represented by $\tau$'s, can coexist despite having the same preference order on the two supplied resources. The monoculture steady state of species $\alpha$ (purple ZNGI) corresponds to the purple point, a state where species $\beta$ (red ZNGI) can successfully invade, and vice versa. The steady state of coexistence is at the black point $X$, where both ZNGIs intersect.

In Fig. S27 one can see that when the environment is shaped by the monoculture $B_\alpha$ is invadable by $B_\beta$ if the depletion time $T_1^{(\alpha)}$ is less than that required for coexistence. On the other hand, when the monoculture $B_\beta$ shapes the environment, the microbe $B_\alpha$ can invade if the depletion time $T_1^{(\beta)}$ is greater than that of coexistence. We get

$$T_1^{(\alpha)} < \frac{(g_{\beta 2} - g_{\alpha 2})\,\log D + g_{\beta 2}\, g_{\alpha 1}\, \tau_{21}^{(\alpha)} - g_{\alpha 2}\, g_{\beta 1}\, \tau_{21}^{(\beta)} + g_{\alpha 2}\, g_{\beta 2}(\tau_{12}^{(\alpha)} - \tau_{12}^{(\beta)})}{g_{\alpha 1}\, g_{\beta 2} - g_{\alpha 2}\, g_{\beta 1}} < T_1^{(\beta)} \qquad \text{(S.92)}$$

From this, (S.84) and (S.88) are easily obtained. Thus here, as usual, one can use either diagrams or more formal approach to derive the criterion.

### K What happens to complementarity and diversity when the growth rate distributions are not similar for different sources

Through the main text we have assumed that the nutritional value of different resources is the same, meaning that the growth rate distributions of the different microbes with respect to different resources are similar to each other. Obviously, if we relax this assumption to a large extent, the conclusions concerning both diversity and complementarity of the mature state community of survivors will require modifications. Yet another conclusion that would have to be revisited in this case is that of the eventual extinction of the anomalous microbes. The lag times that are great equalizers will become more important when the growth rates on different sources can be very different. In this subsection, we will address both diversity and complementarity concerns and then discuss in which way a microbe may benefit from adopting an anomalous metabolic strategy when the distributions of the growth rates on different sources that are available in the environment are not equal.

Suppose there is a universally prized resource on which all the microbes grow comparatively fast, like glucose, and it is present in the environment. We could reasonably expect the distributions of the growth rates on such a resource to have a higher mean value and probably smaller variance than those for a less popular resource. In Fig. S28 a Tilman-inspired diagram (Time plane) shows in which way the difference in the growth rates distributions on two resources $R_1$ and $R_2$ affects the community of three microbes $B_\alpha$ (top choice $R_1$, blue ZNGI), $B_\beta$ (top choice $R_1$, green ZNGI) and $B_\gamma$ (top choice $R_2$, magenta ZNGI). Due to the fact that the growth rates

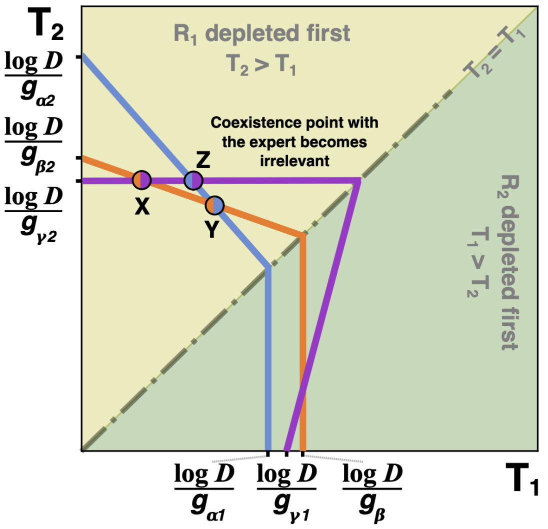

Figure S28: **Complementarity broken by the dissimilarity of the growth rates distributions on two sources.** The ZNGI lines of the experts on $R_1$ (the microbe $B_\alpha$, top choice $R_1$, blue ZNGI) and on $R_2$ (the microbe $B_\gamma$, top choice $R_2$, magenta ZNGI) intersect at the point **Z**, which is driven out of the Pareto-optimal boundary by the segment **XY** of the ZNGI line of a microbe $B_\beta$ (top choice $R_1$, green ZNGI). Thus the experts cannot co-exist in presence of $B_\beta$. The intersection point **X** of the ZNGI of $B_\beta$ (green) and that of $B_\gamma$ (magenta) can be realized at small values of $R_1/R_2 \ll 1$. The intersection point **Y** of the ZNGI of $B_\alpha$ (blue) and that of $B_\beta$ (green) may or may not be realized as a co-existence steady state at a balanced resource supply $R_1/R_2 = 1$, depending on the set of parameters (growth rates, dilution coefficient) of the system.

on $R_1$ are greater than those on $R_2$ any microbe with the top choice $R_2$ (like $B_\gamma$ in our case) will be anomalous by definition. The segment of the Pareto-optimal boundary that allows for the presence of $B_\gamma$ in the resulting uninvadable steady state joins a point at the $T_2$ axis to the point **X**. This means that $B_\gamma$ may be present in the community (always in co-existence with $B_\beta$) only at a very imbalanced nutrient supply, with $R_1/R_2 \ll 1$. This is where we may still have complementarity. The "community of experts" that we have often encountered in our simulations through the main text of the current paper cannot be the resulting set of survivors at the uninvadable steady state. The point **Z** that corresponds to such a situation does not appear on the Pareto-optimal boundary.

#### K.1 The criterion for the presence of the complementarity, lags neglected

Now we proceed to derive a criterion deciding the fate of complementarity at the balanced resource supply in the case of two resources available in the medium, $R_1$ being the one that provides the fastest growth for all (or most) microbes. Let $B_\gamma$ be an expert on $R_2$. It is reasonable to assume that the depletion order in the case under consideration is such that $R_1$ is consumed first. Under this assumption, the only microbe with the top choice $R_2$ that has a chance to survive is the expert – the one that grows on $R_2$ faster than any other microbe. (If such a microbe prefers $R_1$ to $R_2$, there can be no complementarity with that depletion order, as in this case the expert on $R_2$, through a dilution cycle, will use both resources and grow on each of them faster than any other microbe would on $R_2$ only.) Let $B_\beta$ be the microbe whose ZNGI is the first to intersect with the ZNGI of $B_\gamma$ on the left, with the intersection point closest to $T_2$ axis. Coexistence between them is possible if $B_\gamma$ can grow at the point at which $B_\beta$ monoculture is established.

In the steady state of $B_\beta$ monoculture the depletion times are

$$\begin{aligned} T_1^{(\beta)} &= \frac{1}{g_{\beta 1}} \log\left(\frac{D\,R_1 + R_2}{R_1 + R_2}\right), \\ T_2^{(\beta)} &= T_1^{(\beta)} + \frac{1}{g_{\beta 2}} \log\left(\frac{D\,(R_1 + R_2)}{D\,R_1 + R_2}\right), \end{aligned} \tag{S.93}$$

so that, for $B_\gamma$ to be able to grow, we must have

$$g_{\gamma 2}\left[\frac{1}{g_{\beta 1}} \log\left(\frac{D\,R_1 + R_2}{R_1 + R_2}\right) + \frac{1}{g_{\beta 2}} \log\left(\frac{D\,(R_1 + R_2)}{D\,R_1 + R_2}\right)\right] > \log D. \tag{S.94}$$

The above reduces to

$$\frac{g_{\beta 1}\,(g_{\gamma 2} - g_{\beta 2})}{g_{\gamma 2}\,(g_{\beta 1} - g_{\beta 2}} \log D > \log\left(\frac{D\,R_1 + R_2}{R_1 + R_2}\right). \tag{S.95}$$

Introducing a notation

$$x = \frac{g_{\beta 1}\,(g_{\gamma 2} - g_{\beta 2})}{g_{\gamma 2}\,(g_{\beta 1} - g_{\beta 2})}, \tag{S.96}$$

we get from (S.95):

$$\frac{R_1}{R_2} < \frac{D^x - 1}{D - D^x}. \tag{S.97}$$

As we are especially interested in the case with the balanced resource supply $R_1 = R_2$, we get

$$\frac{D+1}{2} < D^x, \tag{S.98}$$

which, with $D \gg 1$, reduces to

$$\boxed{\frac{g_{\beta 2}\,(g_{\beta 1} - g_{\gamma 2})}{g_{\gamma 2}\,(g_{\beta 1} - g_{\beta 2})} < \frac{1}{\log_2 D}} \tag{S.99}$$

When $g_{\beta 1} \gg g_{\gamma 2}$ that would mean huge dispersion for the growth rate distribution on the resource $R_2$, as (S.99) in this case reduces to simply

$$\frac{g_{\beta 2}}{g_{\gamma 2}} < \frac{1}{\log_2 D} \tag{S.100}$$

for a microbe $B_\beta$ with some average (or even greater than average) growth rate on $R_2$.

If however

$$g_{\alpha 1} - g_{\gamma 2} < \sigma_2 \frac{1}{\log_2 D}, \tag{S.101}$$

where $B_{\alpha 1}$ is the expert on $R_2$ and $\sigma_2$ is the dispersion of the growth rate distribution on $R_2$, then for all practical purposes we can have complementarity at the balanced resource supply, as in (S.99) $g_{\beta 1} - g_{\gamma 2} < g_{\alpha 1} - g_{\gamma 2}$ and we could reasonably expect $g_{\beta 1} - g_{\beta 2} = g_{\beta 1} - g_{\gamma 2} + g_{\gamma_2} - g_{\beta 2} \simeq g_{\beta 1} - g_{\gamma 2} + \sigma_2$ to exceed the dispersion $\sigma_2$.

To be more precise we should put in the right side of (S.101) the mathematical expectations of the maximum values of the growhth rates on the corresponding sources, i. e. **the official practical complementarity criterion for the balanced nutrient supply** should be formulated in the following way:

$$\boxed{g_{1max} - g_{2max} < \sigma_2 \frac{1}{\log_2 D}}. \tag{S.102}$$

We should also note that (S.98) (or its detailed version in Eq. (S.99)) must hold for any microbe with the top choice $R_2$, otherwise the complementary coexistence of $B_\gamma$ and $B_\beta$ will be invadable.

##### K.2 When the diauxic lags enter the picture: complementarity preserved and the anomalous microbes getting their chance

Assume that the microbe $B_\beta$ with the top choice $R_1$ experiences have a diauxic lag period $\tau_{12}^{\beta}$ when the resource $R_1$ is depleted, before switching to consumption of the resource $R_2$. (Let the regrowth lags of all the microbes be approximately equal to each other: the differences may play their role either to disrupt or strengthen the complementarity positions, but the diauxic lag is more important for the reason that for the bacteria with the top choice $R_1$ it is manifested while for those with the top choice $R_2$ it does not occur when the depletion order is first $R_1$ then $R_2$.) Then Eq. (S.93) should be modified so that the depletion time $T_2$ in the steady state occupied by $B_\beta$ alone is

$$T_2^{(\beta)} = T_1^{(\beta)} + \tau_{12}^{(\beta)} + \frac{1}{g_{\beta 2}} \log\left(\frac{D\,(R_1 + R_2)}{D\,R_1 + R_2}\right), \tag{S.103}$$

so that $B_\gamma$ with the top choice $R_2$ can invade if

$$g_{\gamma 2}\left[\frac{1}{g_{\beta 1}} \log\left(\frac{D\,R_1 + R_2}{R_1 + R_2}\right) + \tau_{12}^{(\beta)} + \frac{1}{g_{\beta 2}} \log\left(\frac{D\,(R_1 + R_2)}{D\,R_1 + R_2}\right)\right] > \log D, \tag{S.104}$$

which reduces to

$$D^x \exp(y) > \frac{DR_1 + R_2}{R_1 + R_2}, \tag{S.105}$$

where

$$\begin{aligned} x &= \frac{g_{\beta 1}\,(g_{\gamma 2} - g_{\beta 2})}{g_{\gamma_2}\,(g_{\beta 1} - g_{\beta 2})} \\ y &= \frac{g_{\beta 1}\, g_{\beta 2}\, \tau_{12}^{(\beta)}}{g_{\beta 1} - g_{\beta 2}} \end{aligned} \tag{S.106}$$

Note that at large enough lag values $\tau_{12}^{(\beta)}$, when the left-hand side of (S.105) is greater than $D$, Eq. (S.105) is automatically satisfied for all values of $R_1/R_2$. So, we will always have complementarity if

$$\boxed{\tau_{12}^{(\beta)} > \frac{g_{\beta 1} - g_{\gamma 2}}{g_{\beta 1}\, g_{\gamma 2}} \log D} \tag{S.107}$$

for all $B_\beta$ with the top choice $R_1$. If $g_{1\,min} > g_{2\,max}$, we will also have an anomalous microbe among survivors: it will be the expert on $R_2$ among those whose top choice is also $R_2$. This is where **the anomalous metabolic strategy** pays off: when the lag is long enough, all the microbe with the top choice $R_1$ will compete for that particular nutrient, and, among those, there can be only one survivor. In this case, even if a species grows better on $R_1$, when everyone is in favour of $R_1$ for that same reason, it might make a wise decision to switch to the top choice $R_2$.

If however (S.107) does not hold, then (S.105) reduces to

$$\boxed{\frac{R_1}{R_2} < \frac{D^x\, \exp(y) - 1}{D - D^x\, \exp(y)}}, \tag{S.108}$$

with $x$ and $y$ given by (S.106). Substituting $R_1/R_2 = 1$ and assuming $D \gg 1$ we get the condition under which the complementarity is preserved in the case of the **balanced nutrient supply**:

$$\boxed{\tau_{12}^{(\beta)} > \frac{g_{\beta 1} - g_{\gamma 2}}{g_{\beta 1}\, g_{\gamma 2}} \log D - \frac{g_{\beta 1} - g_{\beta 2}}{g_{\beta 1}\, g_{\beta 2}} \log 2} \tag{S.109}$$

#### K.3 In which way non-similarity of the growth rate distributions on different sources affects diversity

Here we consider, again, an environment with two nutrients available for consumption. If all the growth rates on one nutrient, say, $R_1$ are much greater than those on another nutrient, say, $R_2$, the diversity may be out of the question as the lion share of (multiplicative) growth will be done on the most efficient nutrient, the main object of competition between invading microbes. So in this subsection we concentrate on the possibility for an ordinary (non-expert) species with the top choice $R_1$ to coexist with the expert (the one that grows fastest on $R_1$) with the same top choice.

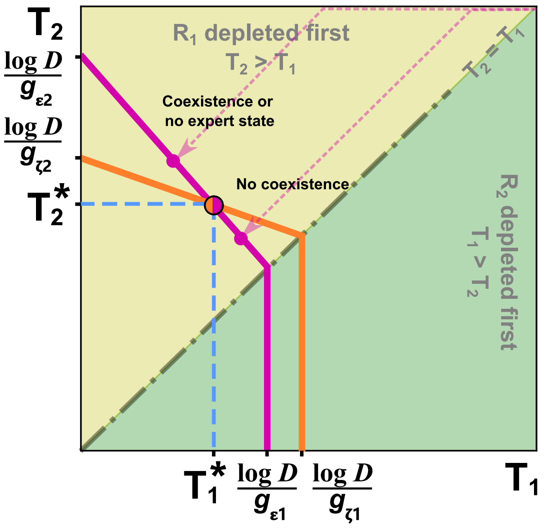

Figure S29: **Diversity in presence of a "super-nutrient"**. An expert microbe $B_\alpha$ (top choice $R_1$, black ZNGI) does not drive out the microbe $B_\beta$ (top choice $R_1$, orange ZNGI) if its one microbe line (black line with an arrow) lands on its ZNGI where the rival species $B_\beta$ can grow. If the opposite is true for any microbe that shares the food preferences with the expert $B_\alpha$, and if the complementarity is not possible, there can be no diversity.

In Fig. S29 the situation favouring coexistence of an expert $B_\alpha$ with another microbe with the same food preferences $B_\beta$ is presented. The one microbe lines of each of two rival species land on the corresponding ZNGI lines (purple ZNGI for $B_\alpha$, orange one for $B_\alpha$) at the point of their monoculture steady states. $B_\beta$ can invade the monoculture of $B_\alpha$ if

$$g_{\beta 1}\, T_1^{(\alpha)} + g_{\beta 2}\, T_2^{(\alpha)} > \log D, \tag{S.110}$$

where

$$\begin{aligned} T_1^{(\alpha)} &= \frac{1}{g_{\alpha 1}} \log\left(\frac{D\, R_1 + R_2}{R_1 + R_2}\right) \\ T_2^{(\alpha)} &= T_1^{(\alpha)} + \frac{1}{g_{\alpha 2}} \log\left(\frac{D\,(R_1 + R_2)}{D\, R_1 + R_2}\right), \end{aligned} \tag{S.111}$$

$T_1^{(\alpha)}$, $T_2^{(\alpha)}$ being the depletion times of the $B_\alpha$−monoculture.

Eq. (S.110) reduces to

$$\boxed{\frac{R_1}{R_2} < \frac{D^h - 1}{D - D^h}}, \tag{S.112}$$

where

$$h = \frac{g_{\alpha 1}\,(g_{\beta 2} - g_{\alpha 2})}{g_{\alpha 1}\, g_{\beta 2} - g_{\alpha 2}\, g_{\beta 2}}. \tag{S.113}$$

In the case of the balanced nutrient supply $R_1 = R_2$ and $D \gg 1$ the condition (S.113) can be reduced to

$$\boxed{\frac{g_{\alpha 2}\,(g_{\alpha 1} - g_{\beta 1})}{g_{\alpha 1}\,g_{\beta 2} - g_{\alpha 2}\,g_{\beta 1}} < \frac{1}{\log_2 D}}. \tag{S.114}$$

If there exists at least one microbe $B_\beta$ such that the condition (S.114) holds, the diversity can be preserved at the balanced resource supply even for non-similar growth rate distributions.

The regrowth lags as well as the diauxic lags may play their part in preserving or destroying the diversity in an obvious way; in the above, we have neglected their effect on the resulting diversity.

##### K.4 Multiple resources, one supernutrient

If there is one supernutrient and a lot of other ordinary nutrients (the latter inspiring similar growth rate distributions for the microbes that use them) present in the environment, we can put together the results of two previous subsections to reach a conclusion concerning the fate of both diversity and the complementarity of a steady state community arising as a result of a long series of species invasions. Indeed, if the parameters of the distributions are such that the complementarity can be preserved in a medium containing one supernutrient and any one of the other ordinary nutrients, we can reasonably expect it to be preserved in the given environment with all the multiple resources present as well. The reasons for that conclusion is that, on average, a microbe uses two or three nutrients during a dilution cycle, and almost never the whole set, as those get consumed side by side, and the multiplication provided by the third resource is even less than that provided by the second one. So the prediction based on two nutrients environment should be good enough. If however the parameters are such that there can be no complementarity, but the diversity is preserved, then we will observe the **complementarity with respect to the second choice**. Indeed, if a microbe $B_\delta$ with the top choice $R_1$ and the second choice $R_3$ can coexist with $B_\alpha$ as well as $B_\beta$ with the top choice $R_1$ and the second choice $R_2$, the niche separation will occur upon depletion of $R_1$, so that they can both survive.

#### References

[1] Manon Barthe, Josué Tchouanti, Pedro Henrique Gomes, Carine Bideaux, Delphine Lestrade, Carl Graham, Jean-Philippe Steyer, Sylvie Meleard, Jérôme Harmand, Nathalie Gorret, et al. Availability of the molecular switch xylr controls phenotypic heterogeneity and lag duration during escherichia coli adaptation from glucose to xylose. *Mbio*, 11(6):e02938–20, 2020.

[2] D Tilman. Resource competition and community structure. *Monographs in population biology*, 17:1–296, 1982.

[3] David Tilman. Resources: A graphical-mechanistic approach to competition and predation. *Amer Nat*, 116(3):362–393, 1980.

[4] D Tilman. Resource competition and community structure. *Monographs in population biology*, 17:1–296, 1982.

[5] Akshit Goyal, Veronika Dubinkina, and Sergei Maslov. Multiple stable states in microbial communities explained by the stable marriage problem. *The ISME journal*, 12(12):2823–2834, 2018.

[6] Veronika Dubinkina, Yulia Fridman, Parth Pratim Pandey, and Sergei Maslov. Multistability and regime shifts in microbial communities explained by competition for essential nutrients. *Elife*, 8:e49720, 2019.

[7] Yuri I Wolf, Mikhail I Katsnelson, and Eugene V Koonin. Physical foundations of biological complexity. *Proceedings of the National Academy of Sciences*, 115(37):E8678–E8687, 2018.

[8] Jef Huisman and Franz J. Weissing. Biological conditions for oscillations and chaos generated by multispecies competition. *Ecology*, 82(10):2282–2695, 2001.

[9] Markus Basan, Tomoya Honda, Dimitris Christodoulou, Manuel Hörl, Yu-Fang Chang, Emanuele Leoncini, Avik Mukherjee, Hiroyuki Okano, Brian R Taylor, Josh M Silverman, et al. A universal trade-off between growth and lag in fluctuating environments. *Nature*, 584(7821):470–474, 2020.

[10] Jue Wang, Esha Atolia, Bo Hua, Yonatan Savir, Renan Escalante-Chong, and Michael Springer. Natural variation in preparation for nutrient depletion reveals a cost–benefit tradeoff. *PLoS Biol*, 13(1):e1002041, 2015.

[11] Matthew D Rolfe, Christopher J Rice, Sacha Lucchini, Carmen Pin, Arthur Thompson, Andrew DS Cameron, Mark Alston, Michael F Stringer, Roy P Betts, József Baranyi, et al. Lag phase is a distinct growth phase that prepares bacteria for exponential growth and involves transient metal accumulation. *Journal of bacteriology*, 194(3):686–701, 2012.
